# BLink-seq delivers population-scale haplotypes without long reads: a scalable framework for non-model genomics

**DOI:** 10.64898/2026.08.03.742036

**Authors:** Azwad R Iqbal, Pavel V Dimens, Jessica A Rick, Paul R Munn, Adrian J McNairn, Jacob B Landis, Rhiannon Schembri, Yingguang Frank Chan, Marek Kučka, Nina Overgaard Therkildsen, Jennifer K Grenier

**Affiliations:** Section of Natural Resources and the Environment, Ashley School of Global Development and the Environment, Cornell University, Ithaca, NY 14853, USA; Genomics Facility and Innovation Hub, Biotechnology Resource Center, Cornell Institute of Biotechnology, Cornell University, Ithaca, NY 14853, USA; School of Natural Resources and the Environment, University of Arizona, Tucson, AZ 85721, USA; Bioinformatics Facility, Cornell University, Ithaca, NY 14853, USA; School of Natural Sciences, Macquarie University, Sydney, NSW, Australia; Cornell Lab of Ornithology, Cornell University, Ithaca, NY 14850, USA; Groningen Institute for Evolutionary Life Sciences, University of Groningen, Groningen, Netherlands

## Abstract

Information about segregating haplotypes and structural variation (SV) can be extremely rich for a variety of applications in population genomics but remains largely inaccessible for many non-model species. Of the available methods, linked-read sequencing is especially promising for its low cost and scalability, but its adoption remains limited. One existing linked-read method is Haplotagging, which barcodes sequencing reads to reconstruct long molecules that encode haplotype information, with the potential to generate phased whole-genome data and detect structural variants. In this study, we present BLink-seq, a novel Haplotagging method that is compatible with standard short-read next-generation sequencing platforms, is locally reproducible with low-cost reagents, and is scalable for high-throughput sample processing. We optimized library preparation parameters, explored their relationship to linked-read library metrics, and validated phasing performance and structural variant detection in two evolutionary extremes: an experimental *Drosophila melanogaster* cross of inbred lines carrying known inversions, and four Atlantic silverside (*Menidia menidia*) parent-offspring trios sourced from highly outbred, wild-caught populations. We then applied our protocol to a cohort of 376 silversides to demonstrate its scalability and potential for SV detection and genotype imputation. Using BLink-seq, we generated chromosome-scale phased blocks and identified known inversions in both validation datasets. We discovered previously uncharacterized structural complexity within a known adaptive inversion on silverside chromosome 11, demonstrating that linked-read data can refine our understanding of SV architecture beyond what short reads alone can resolve. Finally, we provide a user guide for researchers interested in using BLink-seq.

## Introduction

The development of genome-scale technologies and the reduction of their associated costs have rapidly expanded the scope of evolutionary genomic research to include non-model species (Allendorf et al. 2010; Elmer and Meyer 2011; Funk et al. 2012). Next-generation sequencing (NGS) technologies have allowed researchers to generate increasingly large amounts of genetic data, often spanning thousands to millions of single-nucleotide polymorphisms (SNPs) across whole genomes. In non-model species, these datasets have been generated by several different technologies and methods, including restriction site-associated DNA sequencing (RADSeq) (Davey and Blaxter 2010), pooled sequencing of many individuals within a population (Pool-seq), and, more recently, low-coverage whole-genome sequencing (lcWGS) (Lou et al. 2021), and have been applied to questions in molecular ecology, evolution, and conservation management. While SNPs and other genetic data have undoubtedly been important for research programs, these techniques are inherently limited in their ability to reconstruct haplotypes (stretches of linked alleles occupying the same chromosome copy) (Leitwein et al. 2020) and to detect structural variation (Wellenreuther et al. 2019; Ho et al. 2020) at population scales.

For non-model species, incorporating haplotype information can strengthen the ability to detect selective sweeps (Szpiech et al. 2021), characterize local ancestry within the genome, and reconstruct recent demographic history with greater accuracy and fidelity (Leitwein et al. 2020). For species of conservation interest, haplotypes enable the detection of genomic tracts shared among individuals through inheritance from a common ancestor (identity-by-descent, or IBD), which can be used to estimate migration rates, the degree of admixture, and recent effective population sizes (N_e_). IBD tracts can also be used to detect runs of homozygosity (ROH) in the genome, which can be used to infer the degree of inbreeding or population size reduction. Notably, these methods can be applied to a variety of systems and questions, including assessing genetic risk in imperiled populations/species, detecting introgression in migratory species via local ancestry inference, and studying dispersal and admixture, among others.

While haplotype information is extremely powerful for population genetic inference, the ability to accurately phase genomic variants (i.e., assign variants to one chromosome copy or the other) remains limited to species with existing reference panels (e.g., humans) or to projects with sufficient budgets for direct detection with long-read sequencing technologies. While indirect phasing methods using short-read data exist (Browning et al. 2021; Hofmeister et al. 2023), their efficacy is often contingent on sample size, sequencing depth, and the diversity profile of a given species/population. Furthermore, while structural variants have been shown to underlie adaptive trait variation in several non-model species (Joron et al. 2011; Harringmeyer and Hoekstra 2022; Akopyan et al. 2026), they remain relatively under-studied due to the challenges of robust identification with standard short-read sequencing (Stuart et al. 2026).

Recently, novel linked-read sequencing methods have offered a way to directly phase variants by “linking” short fragments based on shared molecular barcode tagging of the molecule of origin. Linked reads can also be used to detect structural variants (SVs) by searching for multiple sets of linked fragments that span candidate rearrangement junctions (Zheng et al. 2016). Until recently, however, linked-read technology had remained expensive or relied on discontinued platforms (e.g., 10x Genomics Chromium Linked-Reads). A new method called Haplotagging (Meier et al. 2021) offers a robust, cost-effective approach to generating population-scale, haplotype-phased genomic datasets in natural populations. However, even Haplotagging has its challenges: it requires extensive initial investment, know-how, and hands-on time to generate the required reagents; the libraries require custom sequencing parameters; and the relatively limited number of published datasets (Meier et al. 2021; Montejo-Kovacevich et al. 2022; Hooper et al. 2024; Orteu et al. 2024; Blumer et al. 2025; Liu et al. 2025; North et al. 2025; Pal et al. 2025) can make it difficult to design a comprehensive project plan.

In this study, we present BLink-seq (**B**ead-**Link**ed sequencing), a variation of the Haplotagging method that leverages a barcode design compatible with standard NGS sequencing parameters and can be reproduced locally using readily available reagents (Fig. 1a). We validate the potential of BLink-seq in two ways. We first show the ability of BLink-seq to robustly phase and detect known inversions, both in *Drosophila melanogaster* F1 progeny from crosses between well-characterized inbred lines and in Atlantic silverside (*Menidia menidia)* parent-offspring trios, despite extreme differences in genomic diversity profiles between species. We then apply our method to 376 Atlantic silversides, representing a population-scale dataset of a wild non-model species, highlighting the potential for genotype imputation, SV detection, and SV characterization (Fig. 1b). To aid users in designing a successful study, we offer a guide to experimental design covering how to tailor the amount of linked-read information generated and how to approach the analysis of phasing and SV detection.

**Figure 1.**
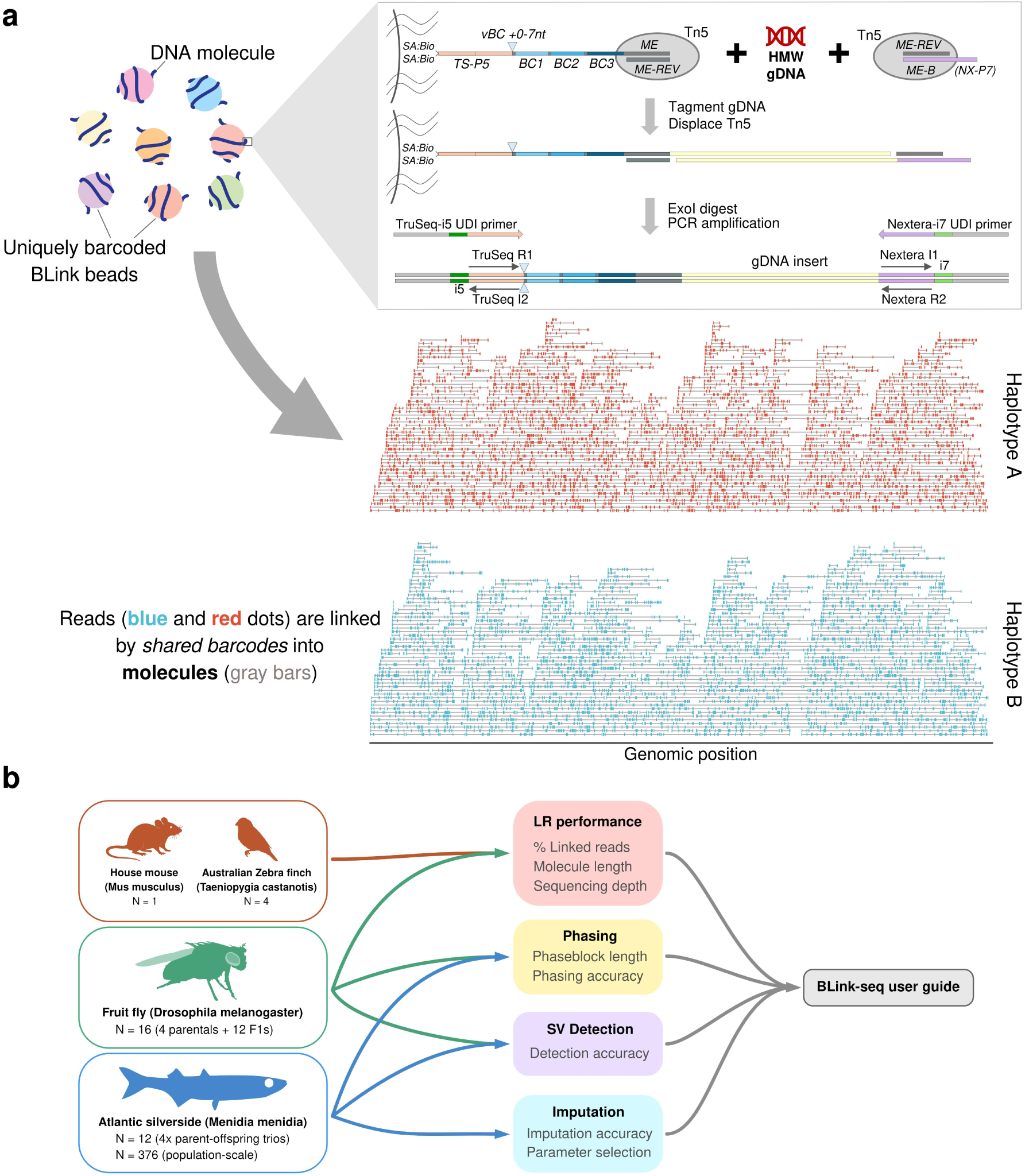
(a) BLink-seq design. BLink beads leverage the tendency of single DNA molecules to wrap around beads; on-bead tagmentation adds a bead-specific barcode to each fragment in the library. Top: BLink bead design with combinatorial barcode, generation of a bead-tethered tagmented fragment, and final amplified library. Components are not shown to scale, and Tn5 is shown as a monomer for simplicity. Each bead has many active transposomes with the same combinatorial barcode, with the potential to generate multiple co-barcoded fragments from a single long gDNA molecule. Bottom: Molecule barcodes can then be used to assign fragments to their molecule of origin, linking reads by shared barcodes into larger molecules (representation inspired by Meier et al. 2021). Molecules can then be assigned to haplotypes using SNP information alongside alignments. (b) Sampling and analytical summary of this study, including species, sample sizes, and analyses completed.

## Methods

### BLink beads

Haplotagging, as described in Meier et al. (2021), is a linked-read sequencing technology that leverages the tendency of single DNA molecules to wrap around microbeads loaded with active, uniquely barcoded Tn5 transposomes, allowing simultaneous tagmentation and co-barcoding of DNA fragments originating from the same molecule. These fragments generate Illumina short-read libraries that capture the molecular barcodes and can be used to “link” co-barcoded reads, thereby reconstructing molecular haplotypes. BLink beads, as used in this project, are individually barcoded streptavidin magnetic beads (Dynabeads™ M-280 Streptavidin, ThermoFisher 11206D) containing a combinatorial barcode and the Tn5 mosaic end (ME) sequence, enabling Tn5 loading and on-bead tagmentation (see Fig. 1a). BLink-seq, unlike the original method developed by Meier et al. (2021), employs a bead-tethered barcoded transposome and an untethered ME-B transposome in solution, allowing easy tuning of the fragment sizes generated during tagmentation by adjusting the amount of ME-B transposome. Our combinatorial barcode design is also unique, employing a variable-length barcode “stagger” segment (1–7 nt, 8 identities) to generate a phased R1 read and three barcode segments (12 nt, 96 identities each), resulting in 7,077,888 possible barcodes. Our method is designed for maximal sequencer compatibility by integrating the combinatorial barcode in-line within the R1 read, which, while reducing effective sequencing coverage of the gDNA fragment, removes the need for custom index read lengths or sequencing primers. BLink beads were prepared through serial split-pool ligation reactions to construct the combinatorial barcodes, and the ME-REV oligo was hybridized to enable Tn5 protein loading to form active transposomes. We have used both Tn5 protein purified in house as well as commercial unloaded Tn5 (Diagenode C01070010). BLink bead performance can be compromised if the Tn5 protein stock contains the N-terminal truncated ‘Inhibitor’ form caused by an alternative initiation codon (Reznikoff 2007); we mutated the methionine at position 56 (M56L) to prevent the ‘Inhibitor’ from being produced (Zhu et al. 2024). A full description of bead assembly and loading is described on the BLink-seq GitHub page (https://github.com/BLinkseq/BLinkseq-Protocol; Iqbal et al. 2026).

### Sample selection for validation and population-scale library preparation

To validate that BLink beads yielded robust and scalable genomic data, we chose to sequence one gDNA sample from a house mouse (*Mus musculus*; C57BL/6 strain) as well as gDNA isolated from individual F1 progeny from a cross between *Drosophila melanogaster* Global Diversity Line strains ZS10 and I13 (Grenier et al. 2015). As both are model species, this allowed for straightforward benchmarking with established genomic resources for validation. The ZS10 × I13 cross was designed specifically because ZS10 is a wild-sourced inbred strain that is a known carrier of large, well-characterized inversions – In(2L)t, In(2R)NS, and In(3R)K – allowing us to directly assess SV detection accuracy. The parental ZS10 strain is heterozygous for all three inversions due to the presence of deleterious alleles and repression of recombination, and thus retains two haplotypes for chromosomes 2L, 2R, and 3R. In contrast, I13 is a homozygous inbred strain not known to carry large inversions. We also explored the effects of DNA degradation on linked-read metrics by sequencing four Australian zebra finch samples (*Taeniopygia castanotis*) representing varying genomic DNA molecular weight profiles (∼4–30 kb mean fragment length) measured on a Femto Pulse instrument (Agilent).

Haplotagging has shown the most promise to enable affordable haplotype-resolved non-model population genomic studies for large cohorts of samples without the need for reference panels. However, there have been relatively few published examples of population-scale Haplotagging datasets due to the method’s relative novelty and limited accessibility. To highlight the potential of BLink-seq for non-model population genomic analysis, we sequenced Atlantic silversides (*Menidia menidia*) generated by laboratory crosses between wild-caught parents. Silversides show one of the highest heterozygosity rates reported for vertebrates (Tigano et al. 2021) and are notable for exhibiting strong local adaptation (Hice et al. 2012) despite pervasive gene flow (Wilder et al. 2020; Akopyan et al. 2026). To first gauge phasing accuracy in this highly outbred species, we sequenced four parent-offspring trios (n = 12 individuals) generated through experimental crosses of wild parents collected from Morehead City, North Carolina, USA (34.72°N, 76.68°W) and Beverly, Massachusetts, USA (42.55°N, 70.90°W), locations that show strong allele frequency differences for multiple inversions linked to latitudinal adaptive divergence (Akopyan et al. 2024; Akopyan et al. 2026). To assess population-scale imputation and SV detection, we sequenced an additional set of 376 F1 offspring from the crossing experiment with wild-caught parents from the two locations.

### DNA extraction

gDNA from the mouse sample was extracted from blood using the Monarch Spin gDNA Extraction Kit (New England Biolabs T3010S). All other gDNA in this study was extracted from tissue samples using a magnetic bead-based protocol (Kučka and Chan 2022) optimized for high molecular weight DNA. For the *Drosophila* dataset, flies were frozen in liquid nitrogen, and genomic DNA was extracted from individual heads to minimize contamination from gut microbiota. For the silversides, DNA was extracted from fin clips that had been preserved frozen (-20°C) in RNAlater for two years.

### Sequencing library preparation

BLink-seq libraries were prepared by incubating sample gDNA with BLink beads and ME-B transposome in the presence of 6.5% PEG8000, 10 mM Mg^2+^ (1× rCutSmart Buffer, New England Biolabs B6004S), and 10% dimethylformamide for 30 minutes in a Thermomixer (Eppendorf) at 55°C with shaking (600 rpm). The tagmentation reaction was then halted and Tn5 displaced by the addition of SDS (final concentration 0.2%), followed by an additional incubation for 7 minutes at 55°C. SDS was quenched by adding Triton X-100 (final concentration 1%), followed by a buffer wash. Beads were then treated with thermolabile Exonuclease I (ExoI, New England Biolabs M0568) for 10 minutes at 37°C to remove excess untagmented barcoded oligos followed by a heat kill for 5 minutes at 65°C and a buffer wash. Depending on the experiment, beads were subsampled to reduce the number of linked fragments (and original molecules) captured in the final Illumina library. Bead-bound tagmented DNA was amplified using 2× High Fidelity PCR Master Mix (New England Biolabs M0541S) and Nextera/TruSeq hybrid UDI primers for 12 cycles before final cleanup, quantification, pooling, and size selection on a PippinHT instrument (Sage Science). For high-throughput (96-sample) batches, we used a MANTIS microfluidic liquid dispenser (Formulatrix) to rapidly dispense master mixes for tagmentation, ExoI treatment, and PCR. See BLink-seq GitHub page (https://github.com/BLinkseq/BLinkseq-Protocol) for full details on BLink-seq library preparation. For a full 96-well plate of samples, library preparation and cleanup can be completed in one day by a single trained individual.

The amount of input DNA and BLink-seq beads can be scaled based on genome size and desired sequencing depth. For each library, we used between 1–1.5 ng of gDNA and 9–22.5 µL of BLink beads (see Table S2 for all library preparation treatments).

### Considerations for sequencing library construction

BLink-seq offers flexibility during library preparation that can alter the properties of the generated data. This includes trade-offs in the amount of linked-read information collected and library complexity, which users can tune to project goals and can affect downstream analysis. To fully leverage linked reads, libraries may need to be sequenced at a higher saturation than for standard whole-genome sequencing (WGS) libraries.

One way to control these trade-offs is to subsample beads after tagmentation and before PCR amplification. By subsampling before amplification, the user can maximize the amount of linked-read information at lower sequencing depths for a smaller set of co-barcoded fragments (and original molecules) per sample. While linked-read information may be maximized by subsampling, the datasets will also reflect lower library complexity with more rapid sequencing saturation. Sample sizes and genome lengths are also important to consider in this paradigm, as the number of individuals sequenced directly affects the sequencing effort allocated per sample on a fixed sequencing budget. These trade-offs are important to consider during experimental design, as certain questions may be better addressed by leveraging linked-read information for fewer molecules per sample across many individuals rather than by sequencing higher complexity libraries at lower depth, or for a smaller cohort of samples.

To highlight the potential and trade-offs associated with bead subsampling, we generated BLink-seq libraries of a single *Drosophila melanogaster* and mouse sample split into technical replicates: one containing the full set of beads used for tagmentation, and three others, representing a subsample of 1/3, 1/6, and 1/12 of the total beads as input to PCR. As sequencing output varied across replicates, we normalized the resulting FASTQ files by subsampling to equivalent reads per library to enable direct comparisons across treatments.

### Data preprocessing and alignment

Preprocessing of raw FASTQs was completed using an in-house pipeline built on cutadapt and Pheniqs (Martin 2011; Galanti et al. 2021) that converts the combinatorial bead barcode into ‘Haplotagging’ format (AxxCxxBxxDxx where the D-segment represents the variable-length barcode) with error correction, and trims 5’ and 3’ adapter sequences. These preprocessing steps have since been incorporated into Harpy, a pipeline for processing linked-read data (Dimens et al. 2025).

Because Harpy is under active development and resolves its dependencies through conda, the specific Harpy release and the versions of the tools it invokes varied over the course of this analysis; we therefore report the tools called by each Harpy module rather than pinned version numbers. Sequences were aligned and duplicates marked with the Harpy Align module using strobealign (Sahlin 2022) and samtools markdup (Danecek et al. 2021) with a deconvolution threshold of 100 kb (-d 100000) and otherwise using default settings to align to the relevant reference genomes (Mouse: GCF_000001635.20, *Drosophila*: GCF_000001215.4, Silverside: GCA_965154125.1, Zebra finch: GCF_048771995.1).

### Single-sample phasing

To assess the phasing performance of BLink-seq, we leveraged both a subset of our *Drosophila* F1 dataset and the silverside parent-offspring trio dataset. For the *Drosophila* dataset, we selected a subset of our F1 libraries with >40% linked reads to ensure that low-quality libraries did not drive phasing results (n = 8 with average depth of 8.6×). Silversides (n = 4 F1s, 8 parents) were sequenced to an average depth of 9.5×.

For both the *Drosophila* and silverside parent-offspring trios, we first performed variant calling on all individuals using GATK 4.1.8 HaplotypeCaller, as implemented in snpArcher (Poplin et al. 2018; Mirchandani et al. 2024). Variant calls were filtered according to GATK best practices and further filtered to remove sites with >75% missingness and minor allele frequencies <0.05 to reduce spurious heterozygous calls that inflate switch error rates during phasing.

Phasing was completed using the Harpy Phase module, which utilizes HAPCUT2 (Edge et al. 2016) with VCF and BAM alignment files as input. HAPCUT2 allows for phasing using only the reads present in the alignment file (treating all reads as “unlinked” short-read fragments) or by also leveraging the barcode tags present in linked-read libraries (“linked”), allowing for the direct comparison of phasing outputs with and without linked-read information.

The Harpy Phase module was run twice for each sample: once with HAPCUT2 using linked-read information with a molecule distance cutoff of 50 kb (-d 50000), and once without linked-read information (-U) with default settings otherwise. For both datasets, we generated a set of pedigree-based (ground-truth) phased variants for benchmarking by using GATK PhaseByTransmission to phase offspring based on the genotypes of parental pairs, supplying a .ped pedigree file alongside variant calls for all members of a parent-offspring trio. We compared these pedigree-based phasing results with those generated by HAPCUT2 for all offspring using whatshap compare (Martin et al. 2016) to compute switch error rates (the proportion of adjacent heterozygous site pairs where the phase relationship differs between the pedigree-based truth and the HAPCUT2 output) in both linked and unlinked HAPCUT2 runs. Phaseblock outputs – contiguous genomic regions phased confidently by the HAPCUT2 algorithm – were sorted and compiled by Harpy for downstream analysis.

### Structural variant detection

Structural variants were called on individual *Drosophila* F1 samples, pooled ZS10 parental *Drosophila* samples, and individual silversides from the parent-offspring trios. SV calls were completed using NAIBR (Elyanow et al. 2018) as implemented in the Harpy SV module using default settings, alongside Wrath (Orteu et al. 2024) using a custom snakemake implementation with 50 kb windows. Wrath Z-score and prediction-band thresholds were set to 3 and 99.5%, respectively. We assessed performance for each program by comparing its ability to detect known inversions segregating within both the *Drosophila* and silverside sample sets. For the *Drosophila* F1 samples, we confirmed the presence of inversions by pre-screening a subset of F1 samples using a PCR assay (Grenier et al. 2015) to detect the presence of inversion breakpoints, alongside further assessment using a principal component analysis on inverted regions to confirm haplotypes for inversions not included in the PCR assay. Silverside inversion haplotypes were confirmed using GTseq calls on SNPs within previously characterized large inversions on chr11, 18, and 24 (Akopyan et al. 2022), while the inversion on chr8 was confirmed via PCA on the inversion region from whole-genome SNPs. While NAIBR also reports duplications and deletions, we focused on inversions as our sample set was designed to test for inversion detection accuracy. An SV call was considered accurate if it bracketed both annotated breakpoints within 5% of the total inversion length (tolerance 0.24–0.72 Mb in *Drosophila*; 0.39–0.47 Mb in silverside).

In addition to single-sample calling, we performed SV calling on samples pooled by inversion haplotypes in both our *Drosophila* F1 and population-scale silverside datasets using Wrath. While Wrath can be run on single individuals, pooling samples can bolster barcode-sharing signals used by the Wrath algorithm to better characterize SVs and reduce output noise (Orteu et al. 2024). We leveraged validated inversion haplotype calls from GTseq data in our population-scale silverside dataset to identify large inversions on chr11, 18, and 24, calling variants for each possible haplotype in groups of 10 individuals. Pooled runs were conducted using 10 kb windows with Z-score thresholds of 3 and prediction-band thresholds of 99.5% initially, with subsequent runs at 25 kb for heatmap visualizations. We plotted barcode sharing heatmaps using Z-scores by window distance to better highlight putative SVs relative to the default percentage-based plots generated by Wrath. We focused our visualizations of Wrath outputs on In(2L)t in our *Drosophila* dataset and the large inversion on silverside chr24 to highlight signals associated with single large “clean” inversions without internal rearrangements or structural complexity.

In the case of the inversion on chr11, we also calculated F_ST_ in 10 kb windows with VCFtools 0.1.16 (Danecek et al. 2011) within the inverted region between heterozygotes (n = 30, as there were no homozygotes for the inverted arrangement in our sample set) and a random sample of homozygous reference individuals (n = 30) to better identify patterns of divergence within the inversion, as our initial Wrath calls suggested possible complexity or nested rearrangements within the larger inversion region.

The properties of linked reads can be leveraged to identify and characterize inversion breakpoints. Inversion breakpoints are expected to show local declines in molecule coverage (see Box 1 in the Supplementary Materials for a definition of “molecule coverage”) for heterozygotes and homozygote carriers. Since molecules are called based on genomic proximity, molecules originating from an inverted haplotype are expected to be truncated at the inversion breakpoint, creating local patterns of low molecule coverage. This decline is expected to be especially pronounced for individuals carrying two copies of the inverted haplotype, with heterozygotes exhibiting an intermediate pattern between the two homozygous arrangements. We used the molecule_coverage.py script in Harpy to calculate molecule coverage in 5 kb windows across the genome to identify regions of low molecule coverage at inversion breakpoints, and normalized coverage patterns relative to genome-wide averages.

### Genotype imputation

Genome-wide variant calls can be a powerful tool for population genomic analyses, but high-confidence variant calling often requires high resequencing depths (>15× coverage) that can be costly and infeasible for population-scale datasets (Watowich et al. 2025). While lcWGS can enable population genomic analyses at scale via genotype-likelihood-based methods (e.g., ANGSD; Korneliussen et al. 2014), many programs require hard-called genotypes and have varying tolerance for the degree of missing data inherent to low coverages (Lou et al. 2021). Genotype imputation can infer missing genotypes either using reference panels of individuals sequenced at high depth (Watowich et al. 2025) or based on variant calls or aligned reads without a reference panel (Davies et al. 2016; Gurke and Mayer 2024). Reference panels can be costly to generate and subject to sampling bias that can undermine imputation accuracy (Zhou et al. 2026). This can be especially challenging in non-model species, where robust reference panel selection often requires information on population structure and demographic histories that may not be readily available.

Linked-read information can be leveraged by STITCH (Davies et al. 2016), an imputation program that does not require a reference panel, to improve imputation accuracy by using linked-read information to inform its underlying haplotype model. STITCH has been utilized in several studies that have used Haplotagging (Meier et al. 2021; Hooper et al. 2024; Pal et al. 2025), though simulations have suggested that incorporating linked-read information only improves imputation when linkage disequilibrium (LD) blocks exceed 1 kb (Meier et al. 2021). We explored two aspects of imputation on BLink-seq datasets: the effect of including BX-tag (linked-read) information on STITCH imputation accuracy and recall, and the effect of imputation and subsequent filtering on phasing accuracy.

To assess imputation accuracy with and without incorporating BX-tag information, we first called variants on our population-scale silverside dataset using GATK 4.1.8 HaplotypeCaller, as implemented in snpArcher. We utilized an existing GTseq dataset (4,243 genome-wide SNPs, see Supplementary methods for details), including the same individuals present in our BLink-seq sample cohort, as ground-truth variant calls to measure accuracy. We performed imputation only on GTseq sites by filtering our genome-wide VCF with bcftools filter 1.20 (Danecek et al. 2021) and ran the STITCH diploid model multiple times by varying parameters K (K = 10–200) and usebx (TRUE = use linked-read information; FALSE = ignore linked-read information), while keeping parameters s = 1 and ngen = 1000 fixed across all runs. We then assessed imputation accuracy by calculating genotype concordance across multiple classes (homozygous major, homozygous minor, heterozygous) using Python scripts adapted from Hooper et al. (2024), filtering to only include sites with INFO scores >0.4.

To assess the impact of imputation on phasing results, we re-called variants on our population-scale silverside dataset, this time including our 12 parent-offspring trio samples along with four technical replicates of progeny sequenced to lower genome-wide coverage (∼6× vs ∼10× in our original trios). We ran STITCH in 500 kb windows along chromosomes 8, 11, 18, and 24 using K = 100, ngen = 1000, and usebx = FALSE. STITCH imputed genotypes were filtered for INFO scores >0.8 for downstream use. We compared phasing accuracy as before with whatshap compare between our pedigree-phased truth set and HAPCUT2 runs with/without imputation.

## Results

### Library preparation decisions illustrate trade-offs in the amount of linked-read information and library complexity

To characterize linked-read metrics relating to experimental design criteria, we prepared technical replicate libraries from single *Drosophila melanogaster* and mouse gDNA samples using a bead-subsampling strategy after tagmentation. After normalizing for total raw read depth for libraries prepared from 5 µL of BLink bead stock (full prep) as well as 3-fold, 6-fold, and 12-fold subsampled beads (respectively: 1.67 µL, 0.83 µL, and 0.42 µL bead stock), we found that the subsampled libraries captured more linked fragments but had lower library complexity (Fig. 2). Thus, despite containing more linked-read information at a given raw sequencing depth, the bead-subsampled libraries contained fewer usable reads due to elevated duplication rates, with lower genome coverage after deduplication (Fig. 2e, Fig. S1a). We observed a limited effect on molecule N_50_, with subsampled libraries exhibiting a modest increase in molecule size (Fig. 2c).

**Fig. 2.**
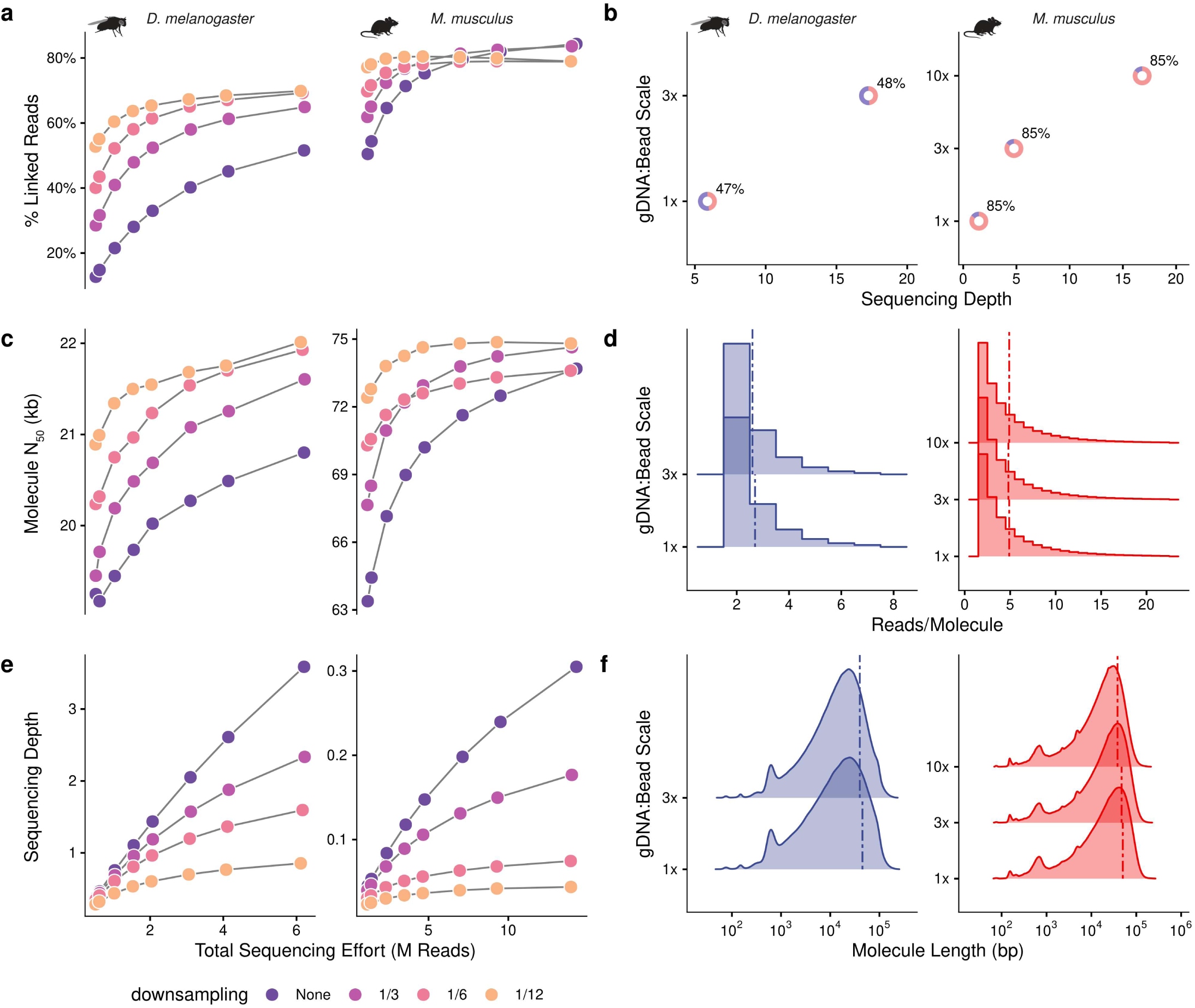
Effects of library preparation variation in key linked-read and general sequencing metrics in *Drosophila* and mouse. (a) Percentage of reads linked by shared barcodes by different treatments. (b) Sequencing depths achieved in different bead/gDNA reaction scales. Donut plots indicate the percentages of reads linked to another by a shared barcode. (c) Molecule N50 by treatment, with “molecules” defined as groups of reads mapped within 100kbp of each other that share a barcode. (d) Distribution of the number of reads within a given molecule by reaction scale. Vertical dashed lines indicate average reads/molecule in a library. (e) Average genome-wide sequencing depth by treatment. (f) Distribution of molecule lengths by reaction scale, with vertical lines representing molecule N50. Comparing the two species datasets, we observed differences in resulting sequencing depths and molecule N50. We attribute the former to differences in reference genome size between species (2.4 Gb for M. musculus versus 143 Mb for D. melanogaster), reflecting the substantially greater number of reads required to reach a given sequencing depth in larger genomes.

When repeating the BLink bead subsampling treatments for a mouse gDNA sample, we observed similar patterns in duplication rates, reads per molecule, and percentage of linked reads increasing with the degree of bead subsampling (Fig. 2a, Fig. S1a,b). Differences in molecule N_50_ recovery, with mouse libraries achieving molecule N_50_ values of ∼73 kb compared to ∼20 kb in *Drosophila* libraries (Fig. 2c), may be driven by differences in the molecular weight of the original gDNA extracts, with more fragmented extracts limiting the maximum size of reconstructed molecules. This is supported by our observation that increasing sequencing effort did not necessarily yield a substantially higher molecule N_50_. The same relationship was evident when comparing zebra finch libraries with varying input gDNA fragment lengths, where libraries generated from higher molecular weight gDNA (longer average fragments) yielded higher molecule N_50_ (Fig. S2). Across these test groups, it is apparent that subsampling beads during library preparation has clear trade-offs with respect to sequencing depth, linked reads, and usable data. For species with relatively small genomes (e.g., *Drosophila*) or for skim-sequencing targeting lower genomic coverage, bead subsampling may be desirable to maximize linked-read information by capturing fewer unique molecules in the dataset. However, for species with larger genomes (e.g., mouse) or when targeting higher molecule coverage, subsampling will generate lower-complexity libraries that rapidly reach sequencing saturation.

To explore how to generate usable libraries for species with larger genomes, we scaled up the initial bead volumes and gDNA input amount during the initial incubation and tagmentation steps, keeping the bead:gDNA ratio the same across replicates, and did not perform any bead subsampling before PCR. Replicates with more beads and gDNA (1×: 1.5 ng of gDNA and 5 µL of beads, 3×: 4.5 ng of gDNA and 15 µL of beads, etc.) yielded higher complexity libraries capturing more barcodes and fragments, and resulted in higher average sequencing depths (*Drosophila*: 5.9× at 1× scale to 17.3× at 3× scale; mouse: 1.5× at 1× scale to 16.8× at 10× scale; Fig. 2b) without substantially elevated PCR duplication rates (Fig. S3) while retaining nearly identical percentages of linked reads (Fig. 2b). Importantly, we observed that increasing the scale of the bead:gDNA incubation did not substantially affect the average number of reads per molecule (Fig. 2d) or molecule N_50_ (Fig. 2f), with the caveat that increasing the scale of the reaction necessitates increased sequencing effort to generate comparable linked-read information.

### Population-scale BLink-seq in a non-model species generates high-quality linked-read data

To demonstrate the scalability of BLink-seq, we generated libraries for 376 Atlantic silverside samples using our high-throughput protocol (see Methods), producing an average of 7.1 ± 2.4 million reads per sample (mean ± SD). After alignment to the *M. menidia* reference genome, samples averaged 2.82 ± 0.96× read depth across assembled chromosomes after marking duplicates, with 341 samples (91%) exceeding 0.5× average depth. We proceeded with all samples in downstream analyses, with the caveat that the lowest-depth samples contribute limited information (four samples produced no usable linked-read data and were excluded from molecule-based statistics).

While per-sample read coverage was relatively modest, linked-read information provided substantially greater effective coverage when reads were aggregated by molecular barcode. Mean molecule depth across the dataset was 27.5 ± 15.1×, approximately 10-fold higher than raw read depth, reflecting the efficiency with which short reads can be linked into longer molecular haplotypes when sequencing effort is spread across a limited number of unique molecules. On average, each sample yielded 4.6 ± 1.2 million unique molecules distributed across 152,000 ± 26,000 unique barcodes (molecule-to-barcode ratio of 30.2 ± 5.5), indicating that each barcode was sampled ∼30 times on average across the genome. Samples averaged 43.5 ± 16.8% linked reads (median 48.5%) with 317 of 376 samples (84%) exceeding 20% linked reads.

Molecule size distributions were consistent with high-molecular-weight gDNA, with a mean molecule N50 of 25.8 ± 14.4 kb. PCR duplication rates averaged 54.4 ± 6.1% and reflected the relatively low gDNA input (1.5 ng per sample) required to maximize linked-read information per library, combined with high sequencing effort relative to library complexity.

### Linked reads generate robust chromosome-scale phasing

Comparing replicate phasing runs with and without linked-read information across both our *Drosophila* and silverside datasets, we found that phaseblocks using linked reads (“linked”) were approximately 20,000-fold longer in *Drosophila* F1s and 6,000-fold longer in silverside progeny than those using only the sequencing reads without linkage information (“unlinked”) (Phaseblock N50 = 21 Mb linked vs 1 kb unlinked in *Drosophila* F1s, N50 = 13 Mb linked vs 2 kb unlinked in silverside progeny) (Fig. 3a,d). The linked phaseblocks span nearly entire chromosomes, while unlinked phaseblocks represent locally phased regions spanning a handful of paired-end reads. Among *Drosophila* samples, shorter phaseblocks were concentrated towards chromosome ends in heterochromatin regions, while euchromatin regions were characterized by large, contiguous, multi-megabase-spanning phaseblocks (Fig. 3b). A similar pattern emerged in the silverside dataset, with shorter phaseblocks also concentrating at chromosome ends (Fig. 3e). Despite phasing across vastly larger genomic intervals, linked reads showed only a marginal increase in absolute switch error rate (+0.33 percentage points in *Drosophila*, +0.27 percentage points in silverside), which was consistent in direction across all sample × chromosome pairs in both species (paired Wilcoxon test, p < 10⁻⁹) (Fig. 3c,f).

**Fig. 3.**
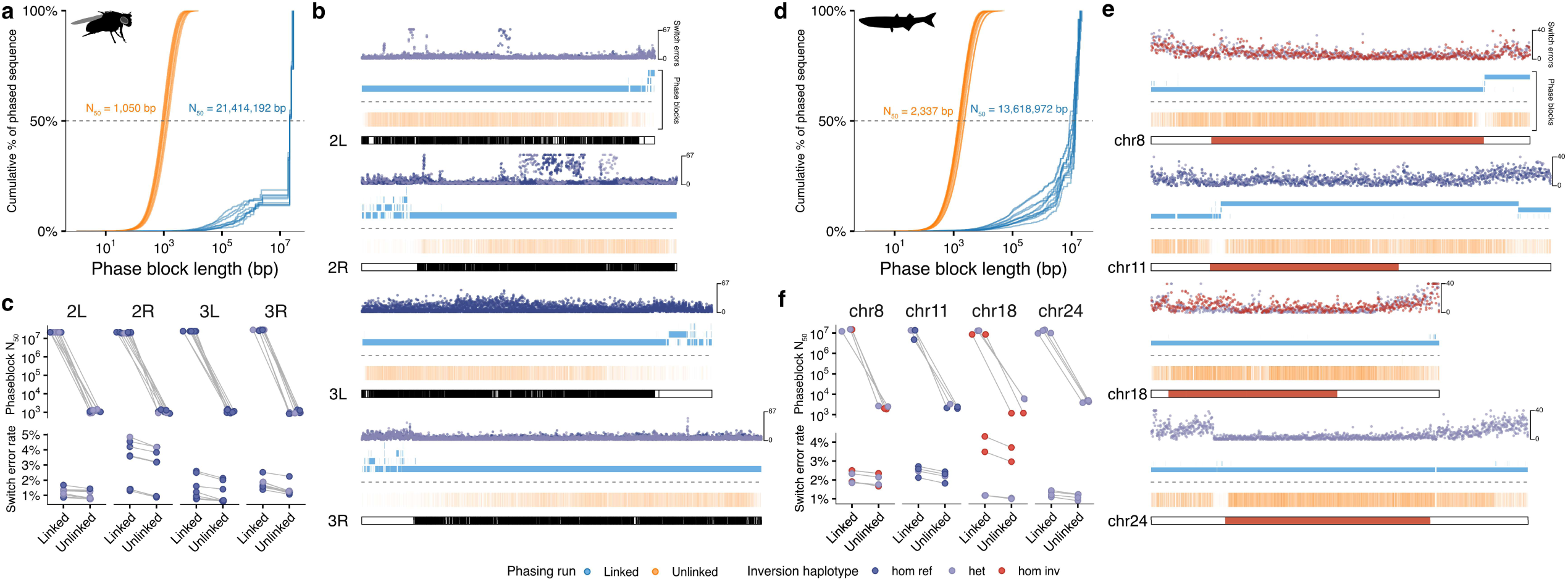
Phasing performance across D. melanogaster and Atlantic silverside (M. menidia) samples. Phasing was completed with (blue) and without b(orange) linked-read information, and we examined switch-error rates and phaseblock statistics comparing inversion haplotypes on chromosomes carrying known inversions. (a,d) Phase block lengths plotted as cumulative percentages of total phased sequences for each sample and species. (b,e) Top: switch error counts of all samples within 50 kb genomic windows colored by sample inversion haplotype. Center: Phase blocks visualized across chromosomes with and without leveraging linked-read information. Bottom: Karyoplots for each focal chromosome. For Drosophila, dark/white regions represent euchromatin/heterochromatin regions, respectively. For silverside, regions in red represent known inversions. (c,f) Phaseblock N50 and switch error rates by sample

### Chromosomal inversions drive patterns in phasing accuracy

As both our *Drosophila* and silverside sample sets contained several large, well-characterized inversions segregating among individuals, we were able to examine the interaction between inversion haplotypes and phasing contiguity and accuracy. Phaseblocks consistently spanned inverted regions in both species, with discontinuities occasionally occurring near inversion breakpoints in silversides (chr8: right breakpoint; chr11: left breakpoint). Notably, switch error rates differed systematically by inversion haplotype – individuals heterozygous for the inversion had consistently lower switch error rates within a chromosome than individuals homozygous for the reference orientation. This is expected as heterozygotes contain long linked tracts of heterozygous variants arising from divergence between the alternate haplotype arrangements, which can then be phased into accurate blocks using linked reads. In contrast, homozygous carriers of the inverted orientation showed elevated switch error rates relative to homozygous reference and heterozygous counterparts. This pattern was present in both linked and unlinked phasing runs, indicating that it was not inherent to a particular mode of phasing, though linked reads appeared to partially amplify this pattern relative to unlinked replicate runs in the same chromosome (Fig. 3f). The overall pattern reflects homozygous inverted individuals carrying fewer heterozygous sites than either heterozygous or homozygous reference individuals, giving the HAPCUT2 phasing algorithm ∼50% fewer sites to phase across the same genomic region.

Imputation recovers phase-informative sites at low coverage

To benchmark STITCH imputation parameters on our population-scale BLink-seq dataset and test whether linked-read information improves imputation in a species with exceptional genome-wide diversity, we imputed genotypes at 4,243 variant sites detected with GTseq across all 376 silverside samples with and without the use of linked-read information and varying the STITCH parameter K. Imputation accuracy was assessed as concordance between imputed genotypes and GTseq ground-truth calls, stratified by genotype class.

Overall genotype concordance was high across all parameter combinations, ranging from 95.9% to 96.6% across sites. Concordance improved with increasing K from 95.9% at K = 10 to a plateau of 96.6% at K = 60–80, beyond which further increases in K provided no additional benefit to accuracy (Fig. S4). The number of sites successfully imputed increased monotonically with K, from ∼39% of GTseq sites at K = 10 to ∼63% at K = 200 (Fig. S5), reflecting STITCH’s conservative handling of sites with limited haplotype support at low K.

Concordance varied substantially across genotype classes. Homozygous major genotypes were imputed with near-perfect accuracy (∼98.4%), while homozygous minor genotypes were moderately accurate (∼87%). Heterozygous sites were substantially more difficult to impute accurately (∼80% mean concordance), with high per-sample variability (SD ∼24%), driven largely by differences in per-sample sequencing depth (Fig. S6). Heterozygous genotypes were consistently imputed poorly in samples below ∼1× depth, whereas samples >2× depth had a mean concordance of ∼90%. This pattern reflects the fundamental constraint that heterozygote calls require both alleles to be sampled with sufficient depth.

Incorporating BX-tag information into imputation did not meaningfully improve accuracy in the silverside dataset. Differences between usebx = TRUE and usebx = FALSE runs were small (<0.35% across all genotype classes and K values) and inconsistent in sign across parameter settings. Similarly, BX-tag inclusion did not meaningfully affect the number of sites successfully imputed (deltas <15 sites out of ∼2,600 imputed). We attribute this to the exceptional genomic diversity and rapid LD decay in silversides (Tigano et al. 2021; Akopyan et al. 2026), which produce LD blocks below the ∼1 kb threshold at which simulations suggest linked-read information improves imputation accuracy (Meier et al. 2021). In species with lower diversity and/or longer LD blocks, linked-read information may provide a meaningful benefit.

Based on these results, we proceeded with chromosome-scale imputation in our merged population-scale and parent-offspring trio dataset using K = 100, ngen = 1000, and usebx = FALSE. To assess whether imputation improves phasing, we compared HAPCUT2 runs on imputed genotypes (INFO > 0.8) against runs on the unimputed GATK calls, using the same pedigree-phased truth set, across the four focal chromosomes in our four trio progeny. In the original trio libraries (∼10× coverage), imputation reduced switch error rates in every sample × chromosome comparison (2.09% to 1.71% on average, a mean reduction of 0.38 percentage points; sign test p = 3.1 × 10⁻⁵) while slightly increasing the number of assessed variant pairs (+2.1%), indicating that the improvement did not come at the cost of phasing fewer sites (Fig. S7a).

In the lower-coverage technical replicates (∼6×), imputation instead raised apparent switch error rates slightly (1.83% to 1.95%), but recovered 18.4% more assessed SNP pairs, indicating imputation recovers phase-informative variants that were initially missing. Comparing arms on a matched number of assessed pairs was more informative: imputed ∼6× libraries yielded 1,336,534 assessed pairs at a mean switch error rate of 1.95%, closely matching the 1,330,837 pairs and 2.09% error rate of unimputed ∼10× libraries (Fig. S7b). Imputation can therefore recover phasing information at low coverage comparable to that obtained from unimputed data at substantially higher depth, though we note that these arms assess overlapping rather than identical site sets.

### Barcode-sharing reliably discovers SVs while revealing structural complexity

We assessed the potential of BLink-seq data for SV detection by comparing SV calls from both NAIBR and Wrath on individual *Drosophila* F1s and a pooled set of ZS10 parental samples (n = 11) and silverside parent-offspring trios (n = 12) carrying known inversions. NAIBR leverages split molecules to call SVs, producing a bedpe file with scored SV calls for downstream analysis. Wrath leverages “barcode-sharing” signals between genomic windows, calling SVs in windows with excess barcode sharing and plotting heatmaps that highlight windows of high barcode sharing indicative of an SV.

Wrath detected every inversion tested in every individual, with the exception of the inversions on silverside chr11 and chr18. The inversion on silverside chr18 is a complex of multiple smaller inversions (Akopyan et al. 2022), and our results may reflect Wrath’s inability to parse a complex signal in single-sample applications. NAIBR was less sensitive overall, missing the chr11 and chr18 inversions entirely, calling only two of eight carriers of the chr24 inversion (25% sensitivity), and missing one of six carriers of *Drosophila* In(3R)K (83% sensitivity). Notably, neither program produced any false positives across the inversions tested (Table S1). In our visualizations of Wrath results for In(2L)t in our pooled *Drosophila* dataset and for the chr24 inversion in our pooled silverside dataset, we identified characteristic signals of excess “barcode sharing” associated with each inversion. *Drosophila* In(2L)t exhibited the expected “bowtie” barcode sharing pattern described in previous literature (Meier et al. 2021; Orteu et al. 2024) while the silverside chr24 inversion exhibited a far more subtle signal, possibly owing to a dearth of molecules spanning the inversion breakpoint (Fig. 4a,b).

**Fig. 4.**
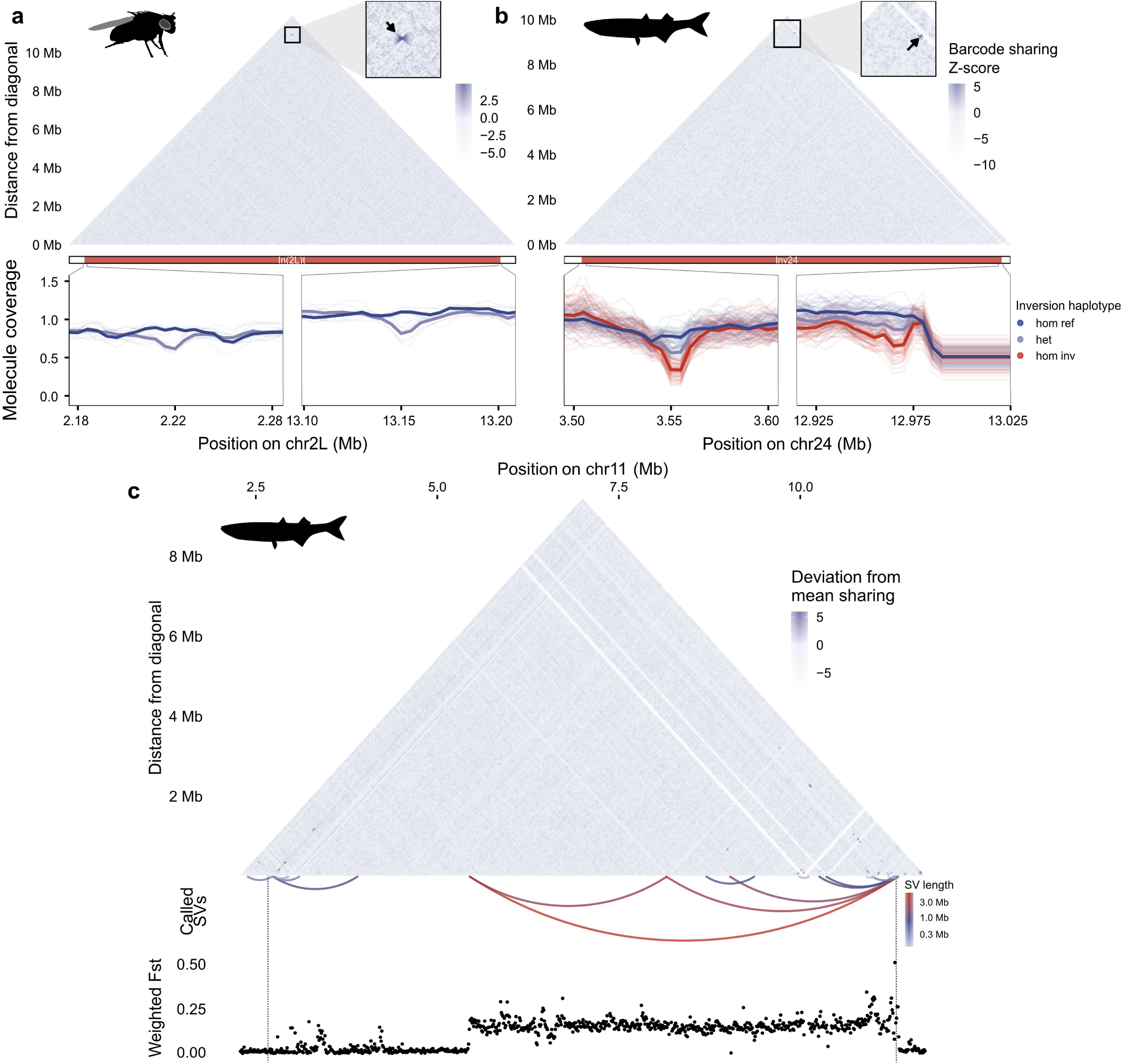
Structural variant detection with linked reads. (a,b) Top: Structural variant detection by comparing barcode-sharing patterns across genomic windows in D. melanogaster and Atlantic silverside. Bottom: Inversion breakpoints are associated with declines in molecule coverage in individuals carrying inverted haplotypes. (c) Examining the canonical inversion region of silverside chr11 (dotted vertical line) reveal patterns of structural complexity associated with elevated differentiation (FST) in heterozygotes vs non-carriers. Top: Barcode-sharing heatmap. Middle: SVs called by Wrath, represented by arcs and colored by SV length. Bottom: FST between inversion carriers and non-carriers in 10 kb windows within the canonical inversion region.

Examining the breakpoints of these inversions more closely, we identified declines in molecule coverage associated with inversion haplotypes (Fig. 4a,b bottom). This was most pronounced in our silverside dataset, as we had a robust sampling of all three inversion haplotypes, but it was also notable in our *Drosophila* dataset, in which all carriers were heterozygous. For the silverside chr24 inversion, we were able to isolate the left breakpoint signal to a region at ∼3.55 Mb, showing the expected pattern of homozygote carriers with the lowest molecule coverage, heterozygotes with intermediate coverage, and homozygous non-carriers with no appreciable decline. The right breakpoint of the chr24 inversion was more ambiguous, with an extended region of reduced molecule coverage associated with carrier haplotypes, adjacent to an extended region of extremely low molecule coverage across all samples.

Our initial comparison of NAIBR and Wrath results on silverside chr11 suggested that Wrath was unable to accurately call the inversion based on previously published ‘canonical’ breakpoints. However, upon examining a Wrath analysis of pooled data from heterozygotes (there were no homozygous carriers among our samples), we found signals of multiple SVs within the bounds of the ‘canonical’ breakpoints (Fig. 4c). The largest of these SVs aligned with a block of elevated F_ST_ between heterozygotes and non-carriers based on SNP data, with multiple smaller SVs also called within the high-F_ST_ region. The left breakpoint region of the chr11 inversion exhibited a cluster of adjacent SV calls, but without a large contiguous high-F_ST_ block. Overall, these results suggest that what was previously identified as a single contiguous inversion may instead be a composite signal of multiple, perhaps even nested, rearrangements.

## Discussion

In this study, we demonstrated multiple implementations of BLink-seq, our modified Haplotagging method, highlighting its ability to impute and phase variants and to call and characterize SVs on multiple scales and biological systems. Early testing on control samples with well-characterized genotypes (*Drosophila*, mouse) demonstrated the performance of our linked-read method and can guide experimental design for the number of molecules, fragments, and reads required for a project. Our population-scale validation datasets included species with vastly different evolutionary and demographic histories, spanning two extremes. On one end, our *Drosophila* dataset represented partially inbred, laboratory stocks of a species with a relatively small genome (143 Mb), while our Atlantic silverside dataset represented a wild outbred species with high standing diversity (Tigano et al. 2021) and a moderate genome size (∼620 Mb). In both systems, BLink-seq achieved robust, chromosome-scale phasing and sensitive SV detection despite relatively low coverage (<10×). In addition to general validation, our method also generated new insights into the architecture of previously characterized SVs by discovering a potentially complex basis for an inversion on chr11 known to underlie adaptive trait variation in Atlantic silversides (Akopyan et al. 2026). Perhaps most importantly, our method is scalable, repeatable, and cost-effective, with a single 96-well plate completed in under one day of laboratory time and costing ∼$3.50/library in materials and reagents, enabling population-scale, haplotype-aware analyses in non-model species for the first time.

### Insights from linked-read data refine our understanding of SV architecture

Structural variants are increasingly understood to underlie adaptive trait architecture across species (Stuart et al. 2026); however, the internal architecture of apparent inversions can be complex and difficult to characterize with traditional short-read-based methods. In silversides, a large inversion on chr11 has been characterized using linkage mapping (Akopyan et al. 2022) and understood to underlie population divergence (Akopyan et al. 2024) and adaptive variation across a clinal gradient using QTL mapping (Akopyan et al. 2026). This inversion has thus far been understood to be a single rearrangement spanning ∼8.6 Mb, with inverted haplotypes more common in the extreme northern range (Wilder et al. 2020). By examining bead-barcode sharing patterns across chr11 with Wrath in our population-scale silverside dataset, we identified multiple, often overlapping SV signals within the canonical inversion breakpoints using a pool of heterozygous individuals. The largest of these SVs coincided with a block of elevated F_ST_ between heterozygotes and non-carriers, and multiple smaller SVs were also identified within the larger SV call and F_ST_ block. Additional SVs were also called adjacent to the left breakpoint, though this did not coincide with a single contiguous signal of differentiation. Overall, this pattern suggests that the chr11 inversion may be structurally complex, comprising multiple nested inversions or a composite element generated by secondary rearrangements after origin. Alternatively, the observed patterns could reflect heterogeneity in recombination within the breakpoint regions, possibly because the source populations are enriched for heterozygotes. Regardless of the exact underlying mechanism(s) shaping these patterns, our finding highlights the value of linked-read information in characterizing inversion structures, while also suggesting that standard short-read-based methods may miss key structural signals that linked reads provide.

### Structural variant detection exhibits software-specific strengths and weaknesses

While our results suggest that Wrath is an accurate tool for SV discovery with the potential to identify complex rearrangements, it is important to consider the associated trade-offs of its approach compared with NAIBR and other SV calling methods. While accurate in our testing, Wrath is computationally expensive owing to its Jaccard matrix-based approach, with runtime scaling with the number of pooled samples and the window sizes relative to the genomic interval(s) being scanned. Wrath outputs can also be noisy for single-sample SV calling using default calling thresholds, as individual samples can carry limited barcode information at smaller window sizes, leading to spurious SV calls. We opted for relatively large windows (50 kb) and stringent Z-score and prediction-band thresholds (3 and 99.5%, respectively) in our single-sample tests to limit noise, though additional tuning or iterative approaches may be required in specific study systems. Pooling samples can improve the resolution of Wrath outputs by enabling smaller window sizes, thereby enabling the detection of smaller SVs. While we were able to pool samples based on known inversion haplotypes, a naive end user without this information may find it difficult to determine pooling strategies. Unlike NAIBR, Wrath does not explicitly define SV classes; the user is expected to parse the size classes and the barcode-sharing heatmap to determine whether an SV may be an inversion, duplication, or another SV class. Ultimately, Wrath and NAIBR are not mutually exclusive. Instead, it may be advisable to run both programs alongside other software not specific to linked reads, such as local PCA (Li and Ralph 2019) or Delly (Rausch et al. 2012), to generate multiple lines of evidence for putative SVs. We did not explicitly test SV genotyping in this study, though tools leveraging linked-read information for inversion genotyping are currently in development (Temperville et al. 2025). Principal component analysis of inversion regions, alongside diagnostic SNP and PCR-based assays, can be alternatives for genotyping inversions with straightforward haplotype structures.

### Inversion haplotype affects phasing accuracy in interpretable ways

Linked-read sequencing has been posited as a means to generate population-scale phased data in non-model species without the need for reference panels (Lutgen et al. 2020; Meier et al. 2021). While reference panels can be powerful when tuned correctly, sample selection, demographic histories, and population structure can all affect downstream phasing results in ways that can be difficult to parse in many non-model contexts. Linked-read sequencing thus provides an avenue for phasing that leverages the molecule information within a given sample rather than external panels, offering a valuable alternative for non-model species if phasing remains accurate.

When examining phasing accuracy across our validation sample set, we found that switch error rates remained low (<5%, Fig. 3) regardless of phasing method, with linked-read runs showing slightly elevated switch error rates but phaseblocks that were >6,000-fold larger. This finding is consistent with previous studies implementing Haplotagging, which found that phaseblock N_50_ values were several orders of magnitude higher when leveraging linked-read information (Meier et al. 2021; Hooper et al. 2024). A factor obscured by comparisons between linked and unlinked phasing runs is that linked runs encompass many more variants per phaseblock than unlinked runs, introducing additional opportunities for switch errors to occur. One advantage of our silverside validation dataset was the ability to study the relationship between inversion haplotypes and switch error rates. Notably, we found that heterozygotes tended to have lower switch error rates on chromosomes carrying inversions, while homozygote carriers exhibited higher switch error rates regardless of whether linked-read information was leveraged. The mechanistic explanation is intuitive for heterozygotes – inversions suppress meiotic recombination (in heterozygotes) and can preserve linkage disequilibrium over long distances, leading to a high density of heterozygous variant sites. Since HAPCUT2 can only phase heterozygous sites, the inverted region presents many input sites in physical linkage for the algorithm to phase accurately. In individuals homozygous for the inverted arrangement, the apparent increase in switch errors may be linked to low heterozygous SNP density within the inverted region, essentially representing the inverse case to heterozygotes. This may not be a universal trend, as the age of the inversion, its evolutionary history, and selection can all affect patterns of diversity within an arrangement. While higher in homozygous carriers in our sample set, the switch error rates reported here do not necessarily preclude key downstream analyses: recent benchmarking of ancestral recombination graph (ARG) inference methods has shown surprising robustness to computational phasing errors, with the caveat that the study tested statistical phasing algorithms (BEAGLE and SHAPEIT) rather than HAPCUT2 (Wang et al. 2025). While outside the scope of this study, reference bias can inflate switch errors depending on the degree of divergence between the focal species and the reference genome; we caution that species lacking a reference genome or with a highly divergent one may experience compounded phasing errors beyond those reported here.

### A BLink-seq user guide

Haplotagging/BLink-seq remains a relatively new method with limited information available to prospective end users on whether the technology may be appropriate for their research questions. While our study and others have highlighted the technology’s flexibility to address a range of topics in population genomics, it remains relatively complex, requiring users to make many key decisions to successfully complete a project. To aid prospective users, we have developed a flowchart that highlights key study design decisions and our recommendations from library preparation through analysis (Fig. 5). In addition, we have developed a user guide containing considerations and recommendations for users in designing and implementing a successful study (Supplementary Materials). While these decisions can be challenging for an end user, we hope this manuscript and accompanying flowchart can be useful when thinking through key study design criteria.

**Figure 5.**
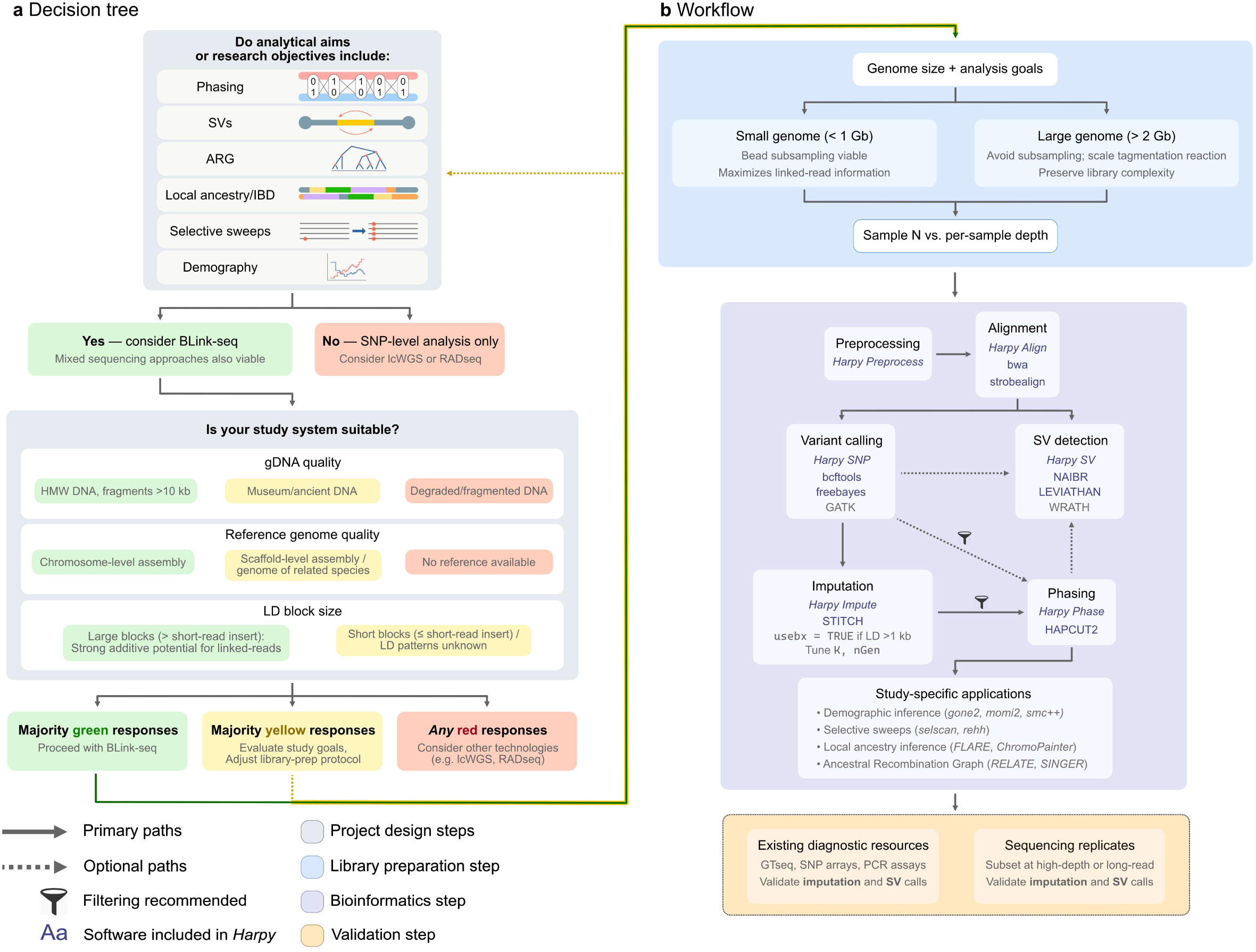
A flow chart to guide prospective users through key decisions and analytical workflows. (a) A decision tree to determine if a study can benefit from BLink-seq, beginning with analytical aims and progressing through key questions regarding study system and available genomic data and resources. For studies that require phased data, are interested in structural variants, or otherwise require haplotype information to address analytical aims, BLink-seq may be suitable. The molecular weight of sample gDNA, alongside the availability of reference genomes and knowledge of LD block sizes can further inform if BLink-seq can enhance a study. (b) Analytical and study design workflows, beginning with library prep and progressing through bioinformatics and validation steps. Example programs and applications are provided for bioinformatics steps, including software included as part of Harpy sub-modules.

### Limitations and future directions

While BLink-seq is a promising technology for non-model species, key methodological and analytical limitations warrant acknowledgment. PCR duplication rates can be higher than those of standard WGS library preparation methods at equivalent sequencing depth, owing to low gDNA input and fragment capture efficiency, which affect library complexity. Sequencing more deeply (with concomitant higher duplication rates) maximizes linked-read information, with a trade-off in the percentage of usable reads. This can be addressed at least in part by tuning bead:gDNA ratios during library preparation, but it remains a key trade-off when considering the value of sequencing read depths vs linked-read information for a given study.

Constructing BLink beads for downstream use requires extensive wet-lab work (typically three days for our protocol), though a single bead prep can scale to thousands of samples depending on library prep parameters. Library preparation itself is similar to tagmentation-based WGS equivalents, and automation with liquid handlers can improve workflow efficiency if available. BLink-seq has not yet been widely applied to museum specimens or other archival samples and may require protocol tuning to account for degraded gDNA that can influence downstream results. BLink-seq also has potential for use in genome assembly and scaffolding akin to Hi-C, though library preparation would similarly need to be modified for this application, for example to reduce barcode ‘clashing’ rates and increase molecule coverage.

Analytically, the software available to fully leverage the potential of linked reads is limited but maturing, with comprehensive pipelines like Harpy (Dimens et al. 2025) providing user-friendly, streamlined workflows compatible with linked-read data. Key areas for enhancement include high-throughput SV genotyping and the development of a linked-read-aware aligner to maintain molecule contiguity and improve alignment across repetitive/unmappable regions. As the adoption of BLink-seq and other Haplotagging technologies increases, we encourage the exchange of knowledge and techniques among research groups to improve existing wet- and dry-lab methodologies.

## Supporting information

Supplemental materials (User guide, supplemental methods, Fig S1-S7)

## Acknowledgments

We would like to thank Harmony Borchardt-Weir for help with DNA extractions, along with Asha Jain, Christine Butler, and Ann Tate for laboratory technical support. The Cornell Office of the Provost provided funding for the Genomics Innovation Hub, which supported the development of BLink-seq. Sequencing services were provided by the BRC Genomics Facility (RRID:SCR_021727) at the Cornell Institute of Biotechnology.

## Data accessibility

Raw whole-genome sequences for this study will be uploaded to NCBI upon peer-reviewed publication of this manuscript. Harpy can be found at https://github.com/pdimens/harpy. The custom fork of Wrath used in this manuscript (“Wrath-plus”) can be found at https://github.com/azwadriqbal/Wrath-plus. Scripts used to assess imputation performance were adapted from https://github.com/dhooper1/Long-tailed-Finch.

## Author contributions

ARI, JKG, and NOT designed the project. JAR performed fieldwork to collect Atlantic silverside specimens, generated GTseq libraries, and processed and analyzed GTseq data. JKG and ARI processed the *Drosophila* samples. JKG, JAR, and RS extracted genomic DNA. JKG designed the BLink beads, and JKG, ARI, and AJM co-developed BLink-seq library preparation protocols, using information provided by YFC and MK. ARI, JKG, PVD, JBL, RS, and JAR generated BLink bead stocks used for this project; AJM purified the Tn5 protein; and JKG, ARI, and RS generated the BLink-seq libraries and whole-genome sequencing data. ARI analyzed the data, with contributions from PVD, PRM, and JKG. NOT and JKG supervised the research. ARI and JKG wrote the manuscript, with edits from all authors.

## Supporting information

Supplementary files are available for download, including the BLink-seq user guide, Figures S1–S7, Tables S1–S2, Supplementary methods, and Box 1.

## References

Akopyan M et al. 2022. Comparative linkage mapping uncovers recombination suppression across massive chromosomal inversions associated with local adaptation in Atlantic silversides. Mol Ecol. 31(12):3323–3341. 10.1111/mec.16472

Akopyan M et al. 2024. Genetic differentiation is constrained to chromosomal inversions and putative centromeres in locally adapted populations with higher gene flow. 2024.10.20.619329 [accessed 2024 Nov 14]. https://www.biorxiv.org/content/10.1101/2024.10.20.619329v1. 10.1101/2024.10.20.619329

Akopyan M et al. 2026. Multiple chromosomal inversions modulate continuous local adaptation along a steep thermal cline. Science. 391(6789):1015–1021. 10.1126/science.ady6774

Allendorf FW, Hohenlohe PA, Luikart G. 2010. Genomics and the future of conservation genetics. Nat Rev Genet. 11(10):697–709. 10.1038/nrg2844

Blumer LM et al. 2025. Introgression dynamics of sex-linked chromosomal inversions shape the Malawi cichlid radiation. Science. 388(6752):eadr9961. 10.1126/science.adr9961

Browning BL, Tian X, Zhou Y, Browning SR. 2021. Fast two-stage phasing of large-scale sequence data. Am J Hum Genet. 108(10):1880–1890. 10.1016/j.ajhg.2021.08.005

Danecek P et al. 2011. The variant call format and VCFtools. Bioinformatics. 27(15):2156–2158. 10.1093/bioinformatics/btr330

Danecek P et al. 2021. Twelve years of SAMtools and BCFtools. GigaScience. 10(2):giab008. 10.1093/gigascience/giab008

Davey JW, Blaxter ML. 2010. RADSeq: next-generation population genetics. Brief Funct Genomics. 9(5– 6):416–423. 10.1093/bfgp/elq031

Davies RW, Flint J, Myers S, Mott R. 2016. Rapid genotype imputation from sequence without reference panels. Nat Genet. 48(8):965–969. 10.1038/ng.3594

Dimens PV et al. 2025. Harpy: a pipeline for processing haplotagging linked-read data. Bioinforma Adv. 5(1):vbaf133. 10.1093/bioadv/vbaf133

Edge P, Bafna V, Bansal V. 2016. HapCUT2: robust and accurate haplotype assembly for diverse sequencing technologies. Genome Res. gr.213462.116. 10.1101/gr.213462.116

Elmer KR, Meyer A. 2011. Adaptation in the age of ecological genomics: insights from parallelism and convergence. Trends Ecol Evol. 26(6):298–306. 10.1016/j.tree.2011.02.008

Elyanow R, Wu H-T, Raphael BJ. 2018. Identifying structural variants using linked-read sequencing data. Bioinformatics. 34(2):353–360. 10.1093/bioinformatics/btx712

Funk WC, McKay JK, Hohenlohe PA, Allendorf FW. 2012. Harnessing genomics for delineating conservation units. Trends Ecol Evol. 27(9):489–496. 10.1016/j.tree.2012.05.012

Galanti L, Shasha D, Gunsalus KC. 2021. Pheniqs 2.0: accurate, high-performance Bayesian decoding and confidence estimation for combinatorial barcode indexing. BMC Bioinformatics. 22(1):359. 10.1186/s12859-021-04267-5

Grenier JK et al. 2015. Global Diversity Lines–A Five-Continent Reference Panel of Sequenced Drosophila melanogaster Strains. G3 GenesGenomesGenetics. 5(4):593–603. 10.1534/g3.114.015883

Gurke M, Mayer F. 2024. GenoPop-Impute: Efficient and accurate whole-genome genotype imputation in non-model species for evolutionary genomic research. [accessed 2026 July 27]. https://www.authorea.com/doi/full/10.22541/au.172515591.10119928/v1. 10.22541/au.172515591.10119928/v1

Harringmeyer OS, Hoekstra HE. 2022. Chromosomal inversion polymorphisms shape the genomic landscape of deer mice. Nat Ecol Evol. 6(12):1965–1979. 10.1038/s41559-022-01890-0

Hice LA, Duffy TA, Munch SB, Conover DO. 2012. Spatial scale and divergent patterns of variation in adapted traits in the ocean. Ecol Lett. 15(6):568–575. 10.1111/j.1461-0248.2012.01769.x

Ho SS, Urban AE, Mills RE. 2020. Structural Variation in the Sequencing Era: Comprehensive Discovery and Integration. Nat Rev Genet. 21(3):171–189. 10.1038/s41576-019-0180-9

Hofmeister RJ, Ribeiro DM, Rubinacci S, Delaneau O. 2023. Accurate rare variant phasing of whole-genome and whole-exome sequencing data in the UK Biobank. Nat Genet. 55(7):1243–1249. 10.1038/s41588-023-01415-w

Hooper DM et al. 2024. Spread of yellow-bill-color alleles favored by selection in the long-tailed finch hybrid system. Curr Biol. 34(23):5444–5456.e8. 10.1016/j.cub.2024.10.019

Iqbal A et al. 2026. BLink-seq Protocol. [accessed 2026 July 31]. https://zenodo.org/records/21723628. 10.5281/zenodo.21723628

Joron M et al. 2011. Chromosomal rearrangements maintain a polymorphic supergene controlling butterfly mimicry. Nature. 477(7363):203–206. 10.1038/nature10341

Korneliussen TS, Albrechtsen A, Nielsen R. 2014. ANGSD: Analysis of Next Generation Sequencing Data. BMC Bioinformatics. 15(1):356. 10.1186/s12859-014-0356-4

Kučka M, Chan YF. 2022. HMW DNA extraction using magnetic beads. [accessed 2024 Nov 14]. https://www.protocols.io/view/hmw-dna-extraction-using-magnetic-beads-b46bqzan

Leitwein M et al. 2020. Using Haplotype Information for Conservation Genomics. Trends Ecol Evol. 35(3):245–258. 10.1016/j.tree.2019.10.012

Li H, Ralph P. 2019. Local PCA Shows How the Effect of Population Structure Differs Along the Genome. Genetics. 211(1):289–304. 10.1534/genetics.118.301747

Liu Z et al. 2025. The fourspine stickleback (Apeltes quadracus) has an XY sex chromosome system with polymorphic inversions on both X and Y chromosomes. PLOS Genet. 21(5):e1011465. 10.1371/journal.pgen.1011465

Lou RN, Jacobs A, Wilder A, Therkildsen NO. 2021. A beginner’s guide to low-coverage whole genome sequencing for population genomics. Preprints. [accessed 2021 Sept 29]. https://www.authorea.com/users/380682/articles/496574-a-beginner-s-guide-to-low-coverage-whole-genome-sequencing-for-population-genomics?commit=a25912ea40ed4aa7f84aeab0b40f1e4d2bea282c. 10.22541/au.160689616.68843086/v3

Lutgen D et al. 2020. Linked-read sequencing enables haplotype-resolved resequencing at population scale. Mol Ecol Resour. 20(5):1311–1322. 10.1111/1755-0998.13192

Martin M. 2011. Cutadapt removes adapter sequences from high-throughput sequencing reads. EMBnet.journal. 17(1):10–12. 10.14806/ej.17.1.200

Martin M et al. 2016. WhatsHap: fast and accurate read-based phasing. 085050 [accessed 2026 July 16]. https://www.biorxiv.org/content/10.1101/085050v2. 10.1101/085050

Meier JI et al. 2021. Haplotype tagging reveals parallel formation of hybrid races in two butterfly species. Proc Natl Acad Sci. 118(25) [accessed 2021 Sept 13]. https://www.pnas.org/content/118/25/e2015005118. 10.1073/pnas.2015005118

Mirchandani CD et al. 2024. A Fast, Reproducible, High-throughput Variant Calling Workflow for Population Genomics. Mol Biol Evol. 41(1):msad270. 10.1093/molbev/msad270

Montejo-Kovacevich G et al. 2022. Repeated genetic adaptation to altitude in two tropical butterflies. Nat Commun. 13(1):4676. 10.1038/s41467-022-32316-x

North HL et al. 2025. The collision of two genomes threatens global food security. 2025.12.25.696198 [accessed 2026 Apr 20]. https://www.biorxiv.org/content/10.64898/2025.12.25.696198v1. 10.64898/2025.12.25.696198

Orteu A et al. 2024. Transposable Element Insertions Are Associated with Batesian Mimicry in the Pantropical Butterfly Hypolimnas misippus. Mol Biol Evol. 41(3):msae041. 10.1093/molbev/msae041

Pal A et al. 2025. Genealogical Analysis of Replicate Flower Colour Hybrid Zones in Antirrhinum. Mol Ecol. 34(22):e70067. 10.1111/mec.70067

Poplin R et al. 2018. Scaling accurate genetic variant discovery to tens of thousands of samples. 201178 [accessed 2026 Apr 21]. https://www.biorxiv.org/content/10.1101/201178v3. 10.1101/201178

Rausch T et al. 2012. DELLY: structural variant discovery by integrated paired-end and split-read analysis. Bioinformatics. 28(18):i333–i339. 10.1093/bioinformatics/bts378

Reznikoff WS. 2007. Tn 5 Transposition. In: Mobile DNA II. John Wiley & Sons, Ltd; p 403–422 [accessed 2026 July 31]. https://onlinelibrary.wiley.com/doi/abs/10.1128/9781555817954.ch18. 10.1128/9781555817954.ch18

Sahlin K. 2022. Strobealign: flexible seed size enables ultra-fast and accurate read alignment. Genome Biol. 23(1):260. 10.1186/s13059-022-02831-7

Stuart KC et al. 2026. A Beginner’s Guide to Structural Variants in Eco-Evolutionary Population Genomics. Mol Ecol. 35(2):e70216. 10.1111/mec.70216

Szpiech ZA, Novak TE, Bailey NP, Stevison LS. 2021. Application of a novel haplotype-based scan for local adaptation to study high-altitude adaptation in rhesus macaques. Evol Lett. 5(4):408–421. 10.1002/evl3.232

Temperville M et al. 2025. SVJedi-Tag : a novel method for genotyping large inversions with linked-read data. In: JOBIM 2025 - Journées Ouvertes Biologie, Informatique et Mathématiques. p 1–9 [accessed 2026 Apr 20]. https://hal.science/hal-05393269

Tigano A et al. 2021. Chromosome-Level Assembly of the Atlantic Silverside Genome Reveals Extreme Levels of Sequence Diversity and Structural Genetic Variation. Genome Biol Evol. 13(6) [accessed 2021 Oct 20]. 10.1093/gbe/evab098. 10.1093/gbe/evab098

Wang L, Deng Y, Nielsen R. 2025. Robustness of Ancestral Recombination Graph Inference Tools to Phasing Errors. 2025.11.24.690249 [accessed 2026 Mar 18]. https://www.biorxiv.org/content/10.1101/2025.11.24.690249v1. 10.1101/2025.11.24.690249

Watowich MM et al. 2025. Best practices for genotype imputation from low-coverage sequencing data in natural populations. Mol Ecol Resour. 25(5):e13854. 10.1111/1755-0998.13854

Wellenreuther M, Mérot C, Berdan E, Bernatchez L. 2019. Going beyond SNPs: The role of structural genomic variants in adaptive evolution and species diversification. Mol Ecol. 28(6):1203–1209. 10.1111/mec.15066

Wilder AP, Palumbi SR, Conover DO, Therkildsen NO. 2020. Footprints of local adaptation span hundreds of linked genes in the Atlantic silverside genome. Evol Lett. 4(5):430–443. 10.1002/evl3.189

Zheng GXY et al. 2016. Haplotyping germline and cancer genomes using high-throughput linked-read sequencing. Nat Biotechnol. 34(3):303–311. 10.1038/nbt.3432

Zhou M, James ME, Engelstädter J, Ortiz-Barrientos D. 2026. Chimeric reference panels for genomic imputation. Genetics. 232(1):iyaf212. 10.1093/genetics/iyaf212

Zhu H et al. 2024. Deciphering gene regulatory programs underlying functionally divergent naïve T cell subsets. 2024.11.06.621737 [accessed 2026 July 31]. https://www.biorxiv.org/content/10.1101/2024.11.06.621737v1. 10.1101/2024.11.06.621737

