## Supplemental materials (User guide, supplemental methods, Fig S1-S7) for "BLink-seq delivers population-scale haplotypes without long reads: a scalable framework for non-model genomics"

#### A BLink-seq user guide

Linked-read sequencing technologies, while powerful, remain relatively underutilized for non-model population genomics. To aid users in designing a successful study, we have developed a brief guide for the decision-making process. In addition, we have provided an overview of how linked-read technology operates, and the associated terminology and metrics associated with LR in **Box 1**.

The first, and perhaps most important, decision is whether linked reads are an appropriate technology for a research project. Haplotype-based analyses, such as local ancestry inference, coalescent-based demographic inference, selective sweep detection, and ARG inference, can benefit from using linked-reads to generate high-quality, chromosome-scale, phased datasets using short-read next-generation sequencing. As we have shown, linked reads can also bolster SV detection and characterization in ways that standard short reads cannot. In contrast, traditional SNP-level analyses, such as population structure inference (ie. STRUCTURE/ADMIXTURE, PCA, NMDS), allele-frequency based summary statistics ($F_{ST}$, $d_{xy}$, pi), or certain demographic inference methods, may not justify the additional costs and analytical overhead that BLink-seq can entail. Ultimately, the question remains whether the study can benefit from information at scales larger than a typical short-read insert. A hybrid approach, utilizing multiple sequencing technologies including linked-reads, has successfully been implemented in several studies exploring structural variation, speciation, and hybridization (Hooper et al. 2024; Blumer et al. 2025; North et al. 2025). Notably, linked-read data can still be used as standard short-read sequencing data for software that cannot utilize linked-read information, offering analytical flexibility.

Beyond analytical considerations, it is also important to consider the study species, genome size, existing genomic resources (i.e., reference genomes, recombination maps, repeat and gene annotations), sample cohort size, and budget available for the project. The availability of a reference genome for the focal species can help reduce spurious variant calls and improve read mapping, ensuring that molecules are accurately reconstructed using bead barcodes within a sample. While often difficult to assess in non-model species, the underlying haplotype structure and linkage-disequilibrium patterns can drive the effectiveness of molecule-based analysis central to linked reads. Long-range molecule information may be uninformative for species with small LD blocks (that is, LD block lengths << molecule lengths), owing to high recombination rates and/or diversity. Sample gDNA quality is also an important consideration, as BLink-seq depends on long molecules wrapping around barcoded microbeads; degraded or fragmented gDNA can reduce molecule size and potentially lead to spurious molecule assemblies due to many random short fragments being captured on a single bead. Museum specimens and historical/ancient DNA, while holding potentially valuable genomic data, are likely challenging for methods like BLink-seq that depend on larger molecule lengths. We therefore recommend high-molecular-weight gDNA with average fragment lengths >10kb as input to maximize the downstream effectiveness of BLink-seq.

Library preparation decisions can have significant implications on downstream results, owing to the interplay between library complexity, genome size, and linked-read information. As we demonstrated in our bead subsampling and reaction scaling experiments, the decision to subsample beads and/or scale the input gDNA mass and bead volumes depends on the study's broader goals and the size of the focal species' genome. For species with larger genomes (>2 Gb – e.g., mammals, some plants), low gDNA inputs or excessive bead subsampling can result in libraries that exhaust complexity before achieving sufficient molecular and sequencing coverage. Alternatively, smaller genomes can allow for additional subsampling or lower DNA input masses to maximize linked-read information per library. Library preparation and sequencing depth should also be tailored to the analytical goals of the project – for studies that primarily focus on SNP-based analyses and/or have smaller sample sizes, sequencing depth may be prioritized at the cost of linked-read information. For studies that rely less on individual genotypes or require robust linked-read information for phasing or SV detection, scaling the input gDNA mass and bead volumes or subsampling beads during library prep can focus sequencing on fewer unique molecules and maximize linked-read information. Notably, imputation via STITCH can help recover and refine variant calls in studies that sequence many individuals at shallow coverage, optionally leveraging barcode tags in its algorithm.

After sequencing, nearly all key analytical steps can be completed with Harpy (Dimens et al. 2025), which includes explicit modules for preprocessing, QC, alignment, SNP calling, SV detection, phasing, and imputation. Harpy automatically generates reports for all modules, which can help assess sequencing quality and analytical outputs for downstream use. While previous Haplotagging studies have used EMA for barcode-aware alignment, we currently do not recommend the software due to unaddressed issues with barcode encoding in the resultant BAMs (see issue #53 at https://github.com/arshajii/ema). Instead, we suggest one of the aligners built into the Harpy Align module, which, while not currently barcode-aware, enables successful, accurate barcode tag encodings and does not interfere with downstream applications. We suggest using both NAIBR (included in the Harpy SV module) and WRATH (installed separately) for SV discovery, for the reasons outlined earlier. LEVIATHAN, another program included in the Harpy SV module, was not tested in this study but can be an additional option. Pooling samples for analysis with WRATH can enhance barcode-sharing signals and enable smaller window sizes to capture smaller SVs, but planning sample pools can be difficult in the absence of known SVs and SV haplotypes. We suggest initially grouping samples by geography/sampling location, or by population cluster if population structure is known. This enables the identification of population-specific SVs and geographic patterns of SV segregation for follow-up.

While variant calling currently does not use linked-read information, genotype imputation via STITCH can optionally leverage barcode tags to construct its internal haplotype models. Previous work using simulations has suggested that barcode tags can benefit species with moderate LD (>1kb block sizes) (Meier et al. 2021), and our imputation results with Silversides suggest that including barcode information offers no meaningful benefit in imputation accuracy, and may actually reduce the number of sites imputed. This is unsurprising as Silversides harbor exceptional levels of genetic diversity and are expected to have small LD blocks outside of known inversions. Phasing via HAPCUT2 can be viable at moderate to low coverage, though long runtimes and high computational requirements can make whole-genome phasing difficult for large datasets or in species harboring high heterozygosity rates. We recommend a targeted approach when relevant, focusing on key chromosomes for analysis rather than full genomes to reduce computational requirements (e.g. the strategy employed by Hooper et al. 2024). Statistical phasing via SHAPEIT5 without a reference panel can be an alternative when sample sizes are large, though it does not currently incorporate linked-read information into its algorithm and is therefore not specific to linked-read datasets.

Phasing and imputation can benefit from subsets of sequencing replicates, either sequenced to high depth or via long-read sequencing, to verify phasing and/or imputation results. Alternatives can include GTSeq (as used in this study) or SNP-array-based datasets that can externally validate imputation results and inform parameter tuning. If within budget and aligned with other project goals, a hybrid approach to sequencing can address limitations between methods and provide opportunities for cross-validation during analysis.

### Box 1 – What are linked reads?

Linked-read data is short read (e.g., Illumina) data, where all the sequences that came from a single unique DNA molecule have the same DNA barcode, which is added during library preparation. This unique quality of sequencing data introduces a few key terms, described here.

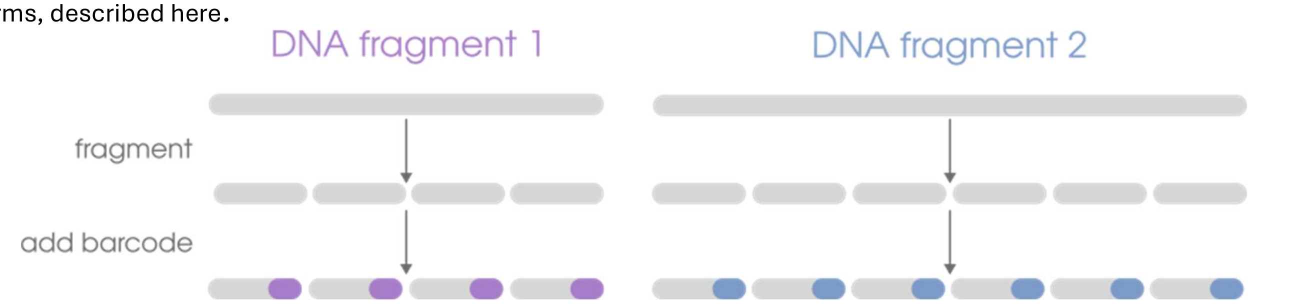


**Linking**
The proportion of sequences sharing barcodes, previously preferred to as "read clouds." If a barcode only appears in one single-end or one paired-end read, it is not "linked" to any other sequences.

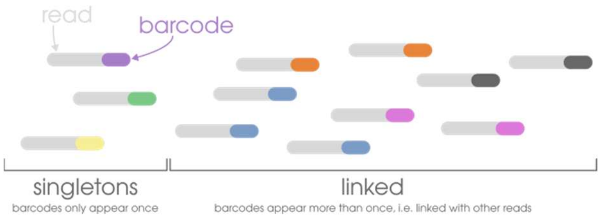


**Molecule Coverage**
The proportion of a unique DNA molecule that is represented by sequences with the same barcode.

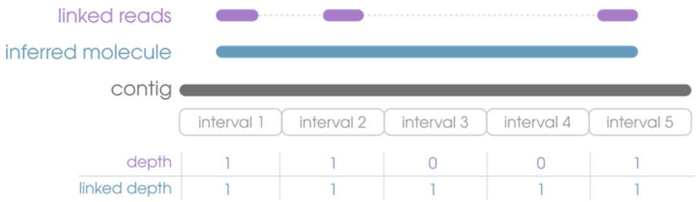


**Linked Depth**
An alternative alignment coverage calculation where all the sequences with a shared barcode are treated as one gapless sequence.

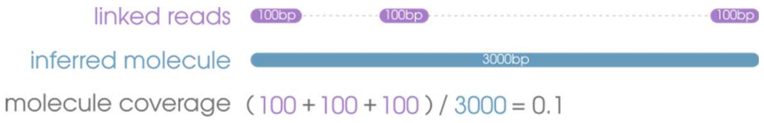


**Deconvolution**
By chance, barcodes can be shared by sequences originating from different molecules ("clashing"). Deconvolution describes the process of determining where clashing is occurring and correcting it by renaming barcodes.

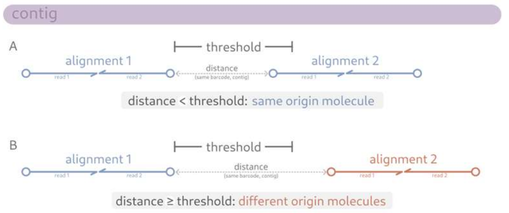

### Supplemental methods

**Atlantic silverside sampling**

In May 2022, adult fish were caught via beach seine at Fort Macon State Park in Morehead City, NC (MCNC), and Obear Park in Beverly, MA (BEMA). Ripe adults were collected in buckets, sexed, and transported to the Rankin Seawater Facility at the University of Connecticut- Avery Point in Groton, CT. At the Rankin Seawater Facility, individuals were strip-spawned and crosses set up in small plastic containers with sheets of wire screen, to serve as substrate for eggs to attach to. For each cross, seawater was placed into the container, followed by female eggs, and finally by male sperm, which was diluted with seawater to facilitate mixing throughout the container. Following strip-spawning, each adult fish was euthanized in an overdose of MS222 and then frozen in individual Ziploc bags. A fin clip was later taken and DNA extracted using a magnetic bead-based extraction protocol (Kučka and Chan 2022).

Following fertilization, screens (with eggs attached) were cut in half and suspended inside of 15 gallon buckets, which were placed within larger tanks, through which water circulated from the seawater facility, held at a constant temperature of 20°C or 26°C. Once larvae started hatching (~115hpf), each bucket was fed an excess of shrimp nauplii for the remainder of the experiment. The offspring in each bucket were thinned at two points during the experiment to ensure equal densities. At an average of 40mm in size (72-79 days post-hatch for 20°C, 55-63 days post-hatch for 26°C), the offspring were phenotyped, euthanized in an overdose of MS222, and then preserved in RNAlater (at room temperature for 5 days and then at -20°C). Fin clips were later taken for DNA extraction using the same magnetic bead-based extraction protocol (Kučka and Chan 2022).

**Genotyping-by-sequencing (GTSeq) of Atlantic silverside samples**

*Primer Design and Testing*

A custom GTseq panel was developed for silversides by GTSeek (Twin Falls, ID). A set of 569 target SNP loci with chromosome and position information were identified as targets for GT-seq genotyping . The chromosome and position information for the target loci were used to pull sequences surrounding the target SNP variants from the Atlantic silverside reference genome assembly for primer design.  A total of 481 primer sets were designed, 406 were chosen for testing, and360 high quality loci retained for the final sequencing panel following an initial round of sequencing.

*Sequencing all individuals*

All parent and juvenile individuals from the experiment were sequenced using this custom silverside GT-seq panel and following Campbell et al. (2015) with modifications. Briefly, primers were pooled and amplified via PCR using the Qiagen Multiplex Plus PCR kit, then diluted 20-fold. Biotinylated barcodes were then added to individual samples in a second PCR step. Following the second PCR, samples were pooled and normalized by using Streptavidin beads. Finally, a bead release PCR step was used to copy bead-bound DNA using bead release primers and Qiagen Multiplex Plus PCR master mix with 6 cycles of PCR. The amplified released product was size-selected for 325-425bp using a two-tailed bead cleanup with Ampure SPRISelect beads (Beckman Coulter Life Sciences). The final product was sequenced on one lane of Illumina NextSeq 2000 (75bp SE) at Novogene. Individuals that did not amplify well or showed evidence of contamination in the first round of sequencing were re-extracted, libraries were prepared, and then those libraries were sequenced on one lane of NextSeq 2000 (75bp PE) at Novogene, following the same methods outlined above.

*Bioinformatic processing and inversion calling*

Raw reads were demultiplexed, followed by adapter trimming using Trimmomatic (v0.39; Bolger et al. 2014) and alignment using bwa-mem2 (v2.2; Li & Durbin 2009) to a subset of the Atlantic silverside reference genome (GCA_965154125.1) containing only the regions targeted in our GTseq panel. Variants were then called using bcftools (v1.20; Danecek et al. 2021) and bamfiles combined for individuals that were replicated within the dataset. Finally, variants were filtered using vcftools (v0.1.16; Danecek et al. 2011) to retain only sites with a maximum of 20% missing data and indels were removed to retain only high-quality SNPs in downstream analyses.

To determine inversion types for known large inversions on chromosome 11, 18, and 24, we performed local PCA using only SNPs located within each of the inversion regions identified in Akopyan et al. (2022) and then classified individuals' inversion types based on those PCA clusters.

**Sample selection for BLink-seq**

We selected a subset of silverside individuals for whole-genome sequencing with BLink-seq. This included four parent-offspring trios (identified using the GTseq data) and 376 additional offspring.

**BLink-seq library preparation**

For the mouse, we used 1.5 ng of gDNA with 10 µL of BLink bead stock. Drosophila F1 and parental libraries were prepared with 1 ng of input gDNA and 9 µL of BLink bead stock, with one exception: a parental ZS10 library was prepared at a scaled-up volume (22.5 µL beads) for downstream comparisons. For the Silverside parent-offspring trios, we used 1.5ng of gDNA with 20 µL of BLink bead stock. To generate our population-scale Silverside dataset, we prepared libraries using a high-throughput version of our library preparation protocol, with 1.5 ng of gDNA and 5 µL of BLink bead stock. See also Table S2.

**References**


Campbell, NR, Harmon, SA, Narum, SR. 2015. Genotyping-in-Thousands by sequencing (GT-seq): A cost effective SNP genotyping method based on custom amplicon sequencing. Mol Ecol Resour. 15: 855-867. https://doi.org/10.1111/1755-0998.12357

Li, H, Durbin, R. 2009. Fast and accurate short read alignment with Burrows–Wheeler transform., Bioinformatics., Volume 25, Issue (14), July 2009, Pages :1754–1760., https://doi.org/10.1093/bioinformatics/btp324

Bolger, AM, Lohse, M, Usadel ,B. 2014. Trimmomatic: a flexible trimmer for Illumina sequence data. Bioinformatics. 2014 Aug 1;30(15):2114-2120. doi: http://doi.org/10.1093/bioinformatics/btu170. Epub 2014 Apr 1. PMID: 24695404; PMCID: PMC4103590.

Danecek, P, et al. 2011. The variant call format and VCFtools. Bioinformatics. 27(15):2156–2158. https://doi.org/10.1093/bioinformatics/btr330

Danecek, P, et al. 2021. Twelve years of SAMtools and BCFtools. GigaScience. 10(2):giab008, https://doi.org/10.1093/gigascience/giab008

Kučka, M, Chan, YF. 2022. HMW DNA extraction using magnetic beads. [accessed 2024 Nov 14]. https://www.protocols.io/view/hmw-dna-extraction-using-magnetic-beads-b46bqzan

#### Supplemental Figures

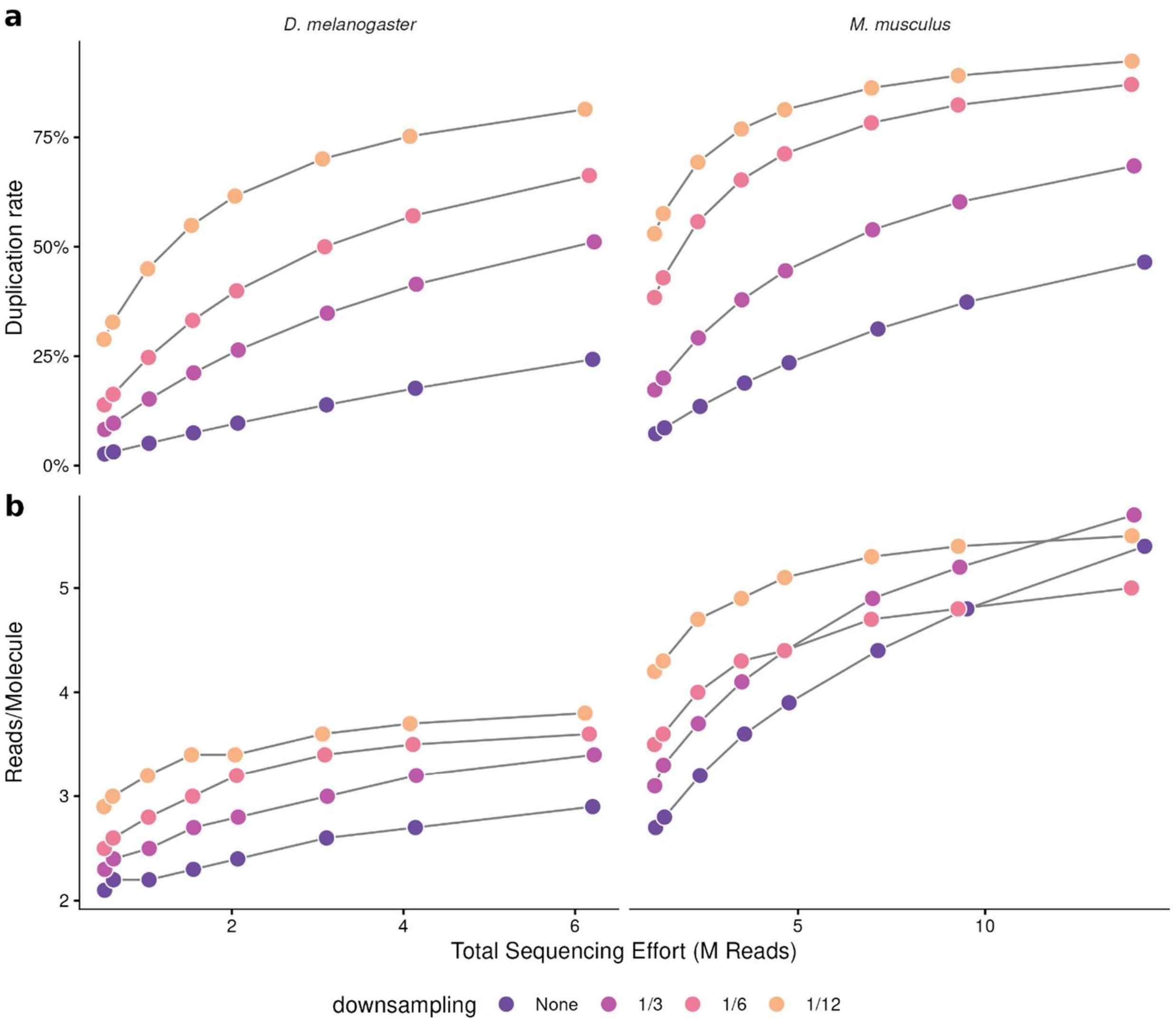


**Fig. S1** (a) Duplication rates by species and bead downsampling (b) Average reads per molecule by species and bead downsampling treatment.

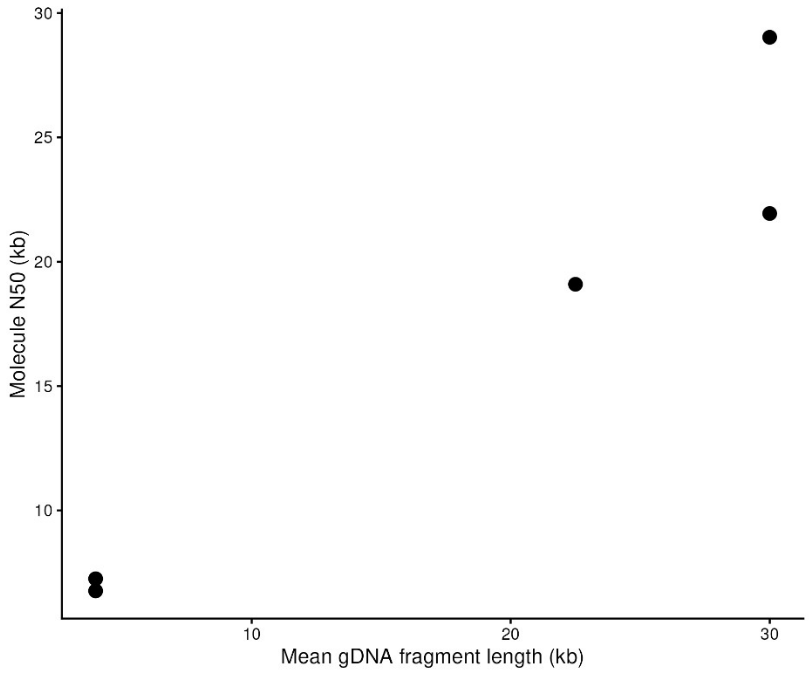


**Fig. S2** Input average gDNA fragment length (molecular weight) vs molecule $N_{50}$ after sequencing and alignment in T. castanotis libraries.

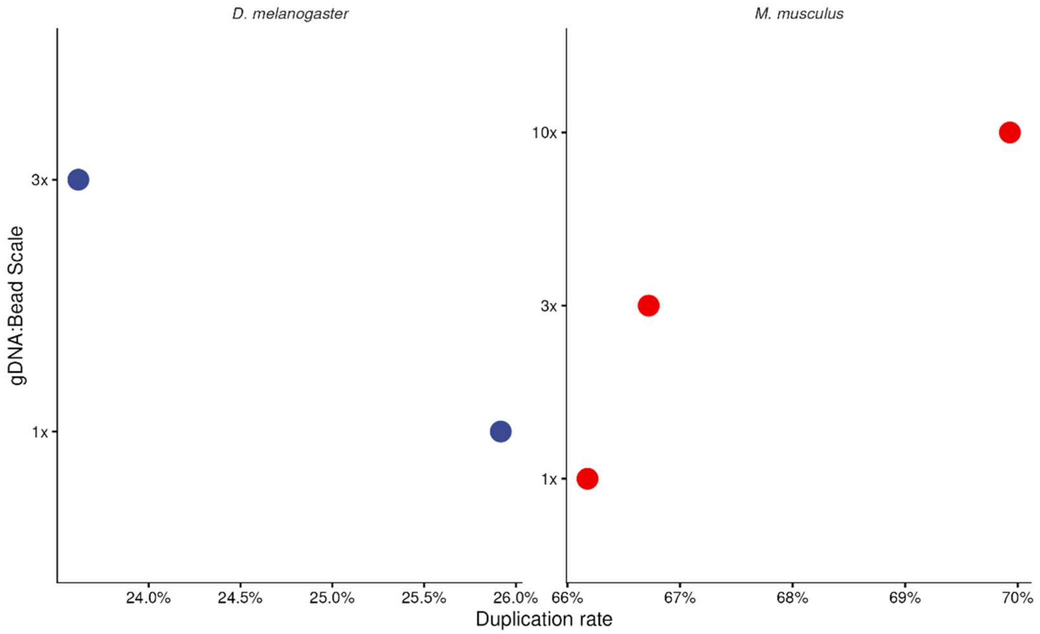


**Fig. S3** Duplication rates by species and gDNA:Bead scale

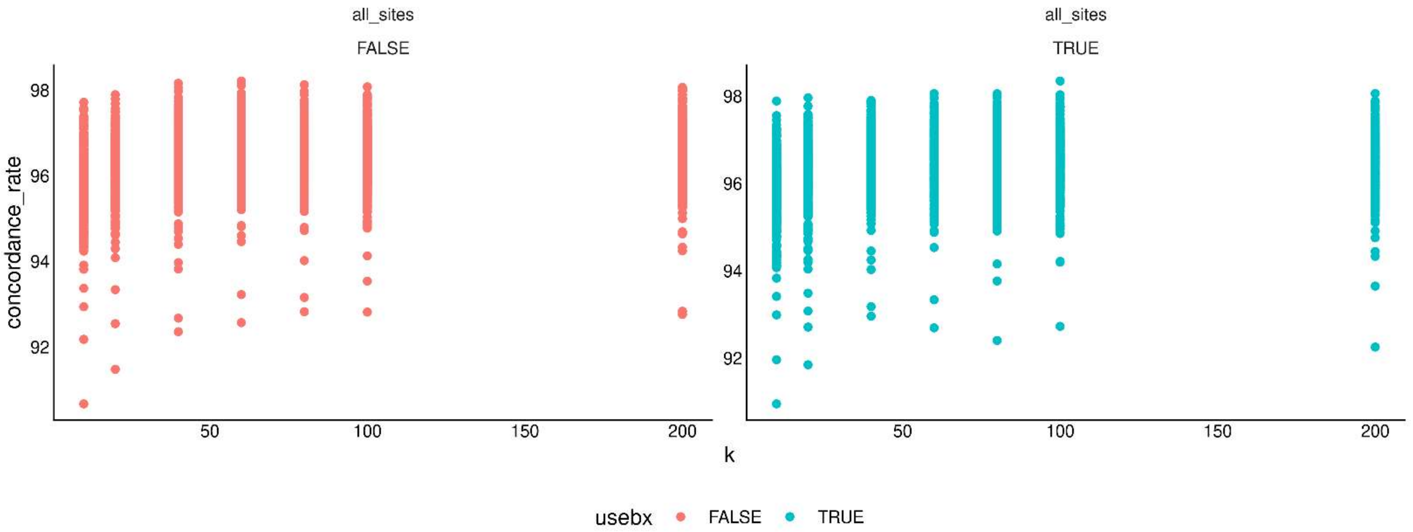


**Fig. S4** Concordance rate of imputed sites across by values of K, colored by whether BX tag information was included (Red = FALSE, Blue = TRUE)

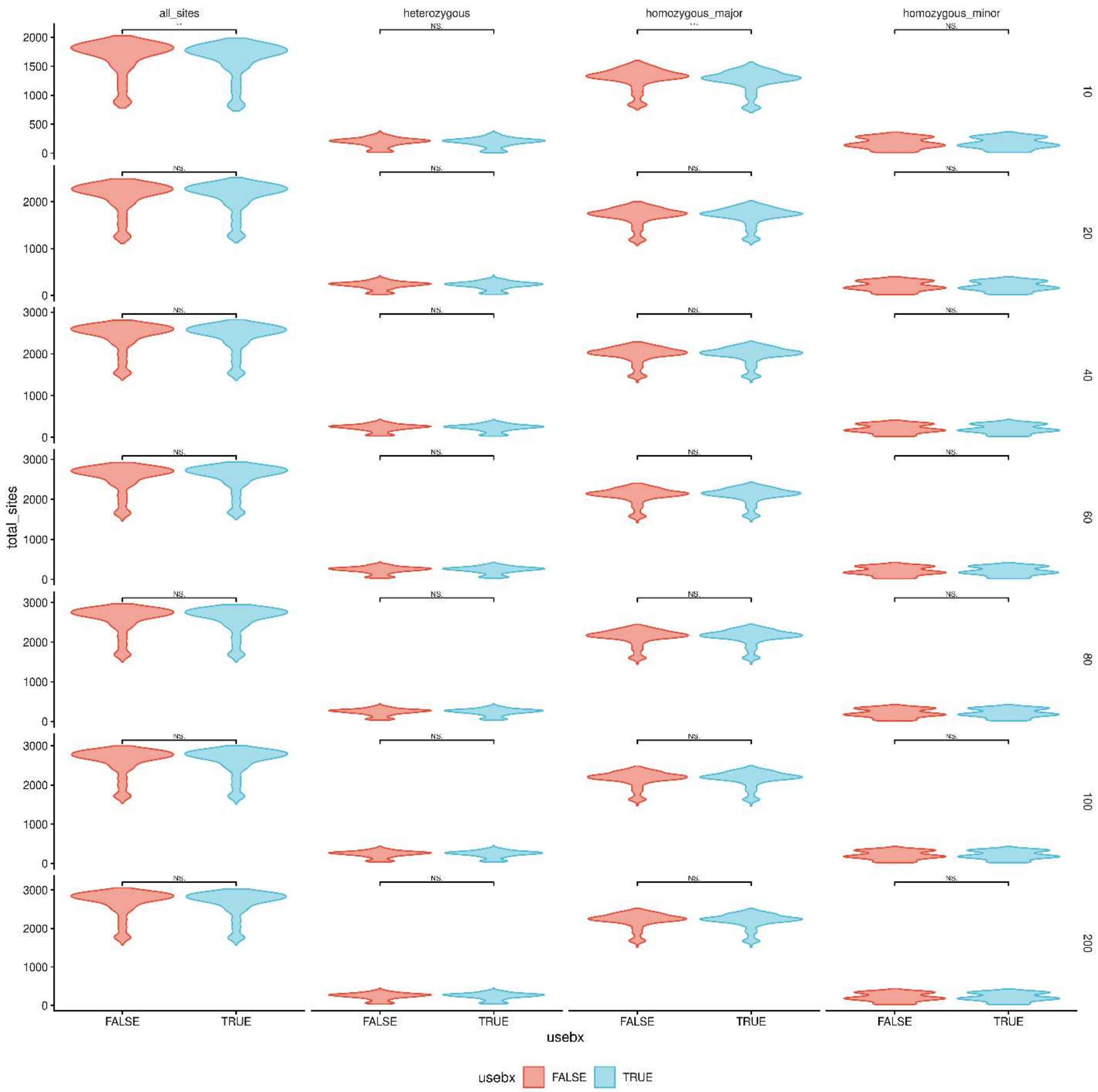


**Fig. S5** Total number of imputed sites across by values of K (right axis, K = 10-200) and stratified by genotype class, colored by whether BX tag information was included (Red = FALSE, Blue = TRUE)

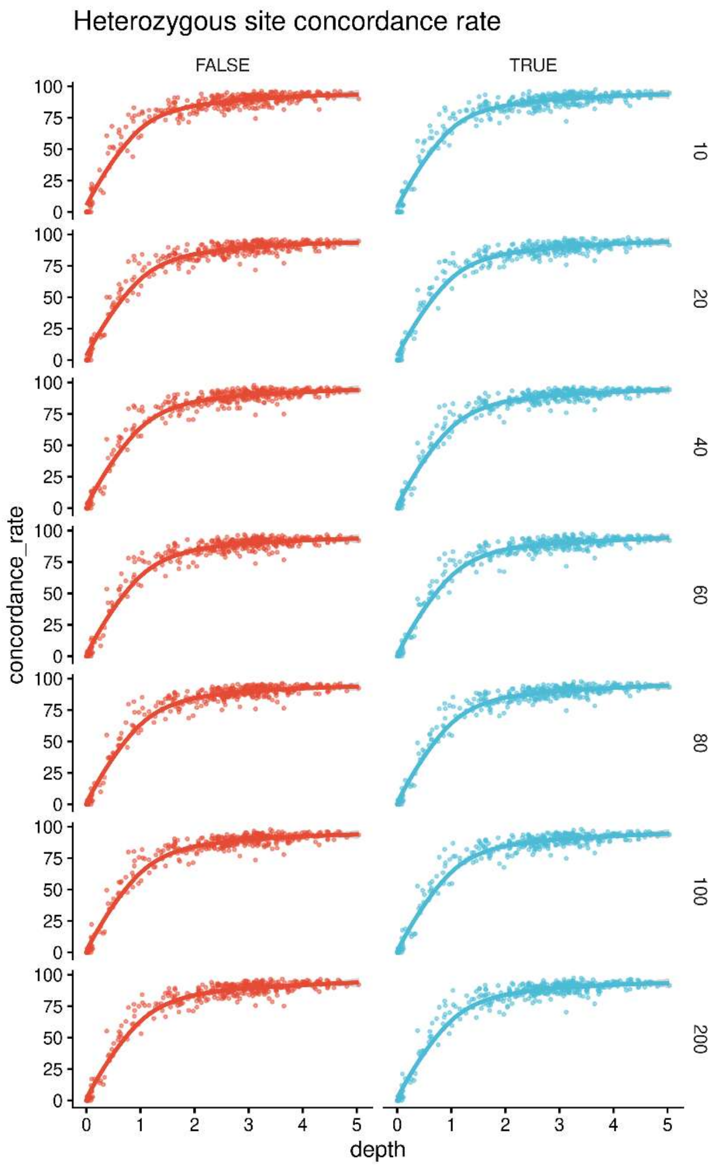


**Fig. S6** Concordance rate of imputed heterozygous sites across by values of K and by average sample sequencing depth, colored by whether BX tag information was included (Red = FALSE, Blue = TRUE)

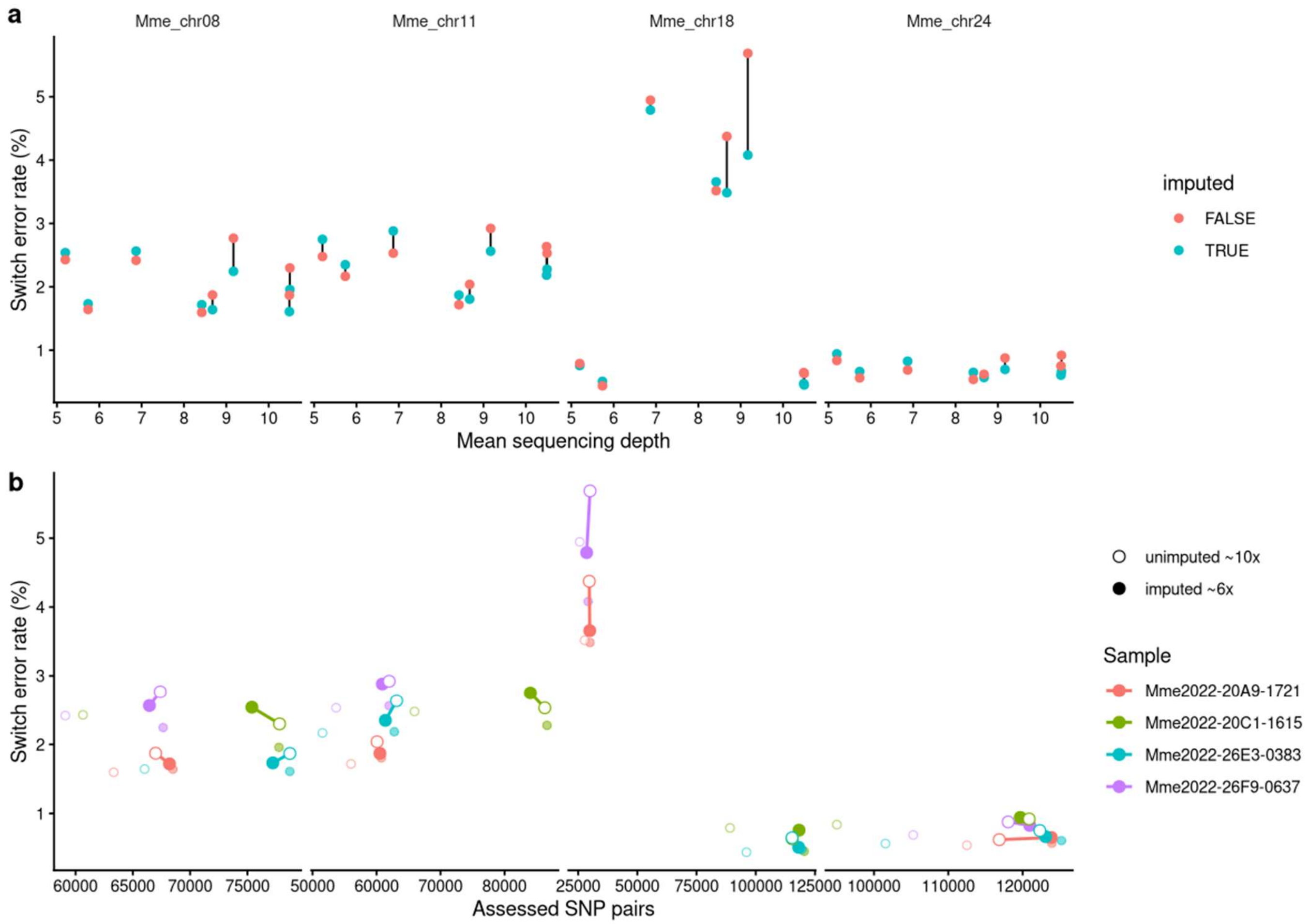


**Fig. S7** Effect of genotype imputation on phasing accuracy in four silverside trio progeny sequenced as high- (~10×) and low-coverage (~6×) replicates. (a) Switch error rate against mean sequencing depth, with lines connecting unimputed and imputed runs of the same library. (b) Switch error rate against the number of assessed SNP pairs, colored by sample. Connected points pair unimputed ~10× with imputed ~6× replicates of the same individual; faded points show the remaining data.

| *Species* | *Inversion* | *Method* | *n* | *TP* | *TN* | *FP* | *FN* | *Sensitivity* | *Specificity* |
|---|---|---|---|---|---|---|---|---|---|
| *D. melanogaster* | In2L | NAIBR | 11 | 7 | 4 | 0 | 0 | 100% | 100% |
| *D. melanogaster* | In2L | WRATH | 11 | 7 | 4 | 0 | 0 | 100% | 100% |
| *D. melanogaster* | In2R | NAIBR | 11 | 3 | 8 | 0 | 0 | 100% | 100% |
| *D. melanogaster* | In2R | WRATH | 11 | 3 | 8 | 0 | 0 | 100% | 100% |
| *D. melanogaster* | In3R | NAIBR | 11 | 5 | 5 | 0 | 1 | 83% | 100% |
| *D. melanogaster* | In3R | WRATH | 11 | 6 | 5 | 0 | 0 | 100% | 100% |
| *M. menidia* | Inv11 | NAIBR | 12 | 0 | 9 | 0 | 3 | 0% | 100% |
| *M. menidia* | Inv11 | WRATH | 12 | 0 | 9 | 0 | 3 | 0% | 100% |
| *M. menidia* | Inv18 | NAIBR | 12 | 0 | 2 | 0 | 10 | 0% | 100% |
| *M. menidia* | Inv18 | WRATH | 12 | 1 | 2 | 0 | 9 | 10% | 100% |
| *M. menidia* | Inv24 | NAIBR | 12 | 2 | 4 | 0 | 6 | 25% | 100% |
| *M. menidia* | Inv24 | WRATH | 12 | 8 | 4 | 0 | 0 | 100% | 100% |

**Table S1**. Inversion detection results from both Atlantic silverside (*M. menidia*) and *Drosophila* samples using both *NAIBR* and *Wrath*. "TP" is a True Positive, "TN" is True Negative, "FP" is False Positive, and "FN" is False negative.
